# Novel dopaminergic neurotransmission in the *Octopus* visual system

**DOI:** 10.1101/2025.04.02.646353

**Authors:** Amy Courtney, Ruth Styfhals, Marie Van Dijck, Jérôme Boulanger, Lieve Geenen, Maxence Lanoizelet, Iris Hardege, Mark Lassnig, Anna M. Jansson, Joseph Gehler-Rahman, Eduardo Almansa, Horst A. Obenhaus, Eve Seuntjens, William R. Schafer

## Abstract

*Octopus vulgaris*, the common octopus, has a complex brain that evolved independently from that of vertebrates, raising the possibility that it may be built using novel circuit motifs or non-canonical neurochemistry. Through systematic characterization of octopus ligand-gated ion channel genes, we identified novel ionotropic receptors for two neurotransmitters, dopamine and acetylcholine, that play key roles in visual circuits. One of these, DopC1, encodes a dopamine-gated cation channel expressed in deeper layers of the optic lobe (inner granular layer and medulla). A second, AChRB1, encodes an acetylcholine-gated anion channel expressed in photoreceptors and dopaminergic outer granular layer neurons. Dopamine drives excitation, particularly in deeper, DopC1 enriched layers, and thereby may play an important role in feed-forward excitation, whereas acetylcholine evokes inhibition, possibly mediating negative feedback from deeper layers. The octopus visual system thus shows fundamental differences in both neurochemistry and wiring compared to mammals, implying distinct mechanisms of visual information processing.

## Introduction

Coleoid cephalopods, such as squid, cuttlefish and octopus, are notable for their large brains and complex behaviour. For example, the nervous system of the common octopus, *Octopus vulgaris*, contains approximately 500 million neurons^1^, more than some mammals^2^, and supports intricate behaviours such as camouflage, tool use, visually-guided hunting, and sophisticated learning and memory^3^. However, octopuses and humans diverged evolutionarily approximately 600 million years ago, early in the bilaterian lineage, from a common ancestor with a much simpler nervous system^4^. Thus, the complex brains of cephalopods appear to have evolved independently from those of vertebrates. Gaining a better understanding of cephalopod neural circuits will therefore shed light on the general principles that underpin complex behaviour across species, as well as discover new biology and species-specific innovations.

The visual system represents a particularly relevant opportunity to compare cephalopod and vertebrate neural circuitry. Vision is an important sensory modality for cephalopods, driving complex visually guided behaviours including camouflage, hunting, mating and learning^5^. Like vertebrates, cephalopods have two camera-type eyes that project visual scenes onto a retina consisting of a single layer of photoreceptors, which direct their output to the cortex of the optic lobe (OL), comprised of a superficial outer (OGL) and deeper inner (IGL) granular layer. These neuronal cell layers are separated by a plexiform, neuropil layer that also includes the end terminals of photoreceptors^6^. The OGL consists exclusively of neurons with amacrine morphology, some of which receive inputs from photoreceptors and extend processes laterally within the plexiform layer. The IGL contains a diverse population of neurons, including amacrine-like cells that arborize in the plexiform layer, centrifugal cells that send projections through the optic nerves back to the retina, and a variety of centripetal cell types that extend processes through the plexiform layer and into the medulla^5^.

The cortical layers of the OL, like the bipolar and granular cell layers in the mammalian retina, are thought to carry out the first stages of visual information processing^7^, and are hence often referred to as the “deep retina”. By way of comparison, in the vertebrate retina, glutamatergic photoreceptors make excitatory retinotopic connections with off-bipolar cells and inhibitory connections with on-bipolar cells; these bipolar cells in turn make excitatory glutamatergic connections with retinal ganglion cells. Laterally-connected horizontal cells (in the outer plexiform layer) and amacrine cells (in the inner plexiform layer) receive excitatory glutamatergic inputs from photoreceptors or bipolar cells, and send inhibitory GABA or glycinergic outputs to laterally-displaced cells in the layer^8^. The vertebrate retina is thus characterized by both excitatory and inhibitory descending signaling, inhibitory lateral signaling, and a lack of ascending feedback. It is unclear whether these microcircuit properties are paralleled in the optic lobes of cephalopods.

To understand how cephalopod optic lobe circuits process information, it is necessary to define the functional properties of their constituent neurons. In octopus, the main neurotransmitters in the optic lobe are glutamate, acetylcholine (ACh) and dopamine^9,10^. While radiochemical experiments previously indicated that acetylcholine and dopamine are synthesized in the retina (eye)^11^, more recent electrophysiological recordings on optic lobe slices from *Octopus vulgaris* indicate that photoreceptors release glutamate as an excitatory neurotransmitter from terminals in the plexiform layer, and that this release is inhibited by acetylcholine^12^. However, although inhibitory neuromodulatory effects of dopamine have been reported in cuttlefish^13^, the role of dopamine in octopus remains unknown. Likewise, while glutamate responses in octopus are likely mediated by AMPA-like receptors^12^, the receptors mediating cholinergic and dopaminergic signaling in the octopus optic lobe have not been identified. Since invertebrate nervous systems can express ionotropic receptors whose ligands and ion selectivity differ markedly from the channels found in vertebrates^14–17^, the octopus brain might in principle use novel ion channels and unexpected neurochemistry for fast neurotransmission.

In this study, we present the first steps toward mapping the pathways of chemical neurotransmission in the *Octopus vulgaris* hatchling. By expressing octopus ion channel genes in *Xenopus* oocytes, we characterised two new ionotropic receptors from the *Octopus* visual system, including the first example of a dopamine-gated cation channel as well as an anionic ACh receptor specifically expressed in dopaminergic neurons. Through functional imaging from paralarval brain slices, we find that dopamine acts as a major excitatory neurotransmitter in the optic lobe, mediating feed-forward signaling from the OGL to deeper layers, while ACh acts as an inhibitory neurotransmitter that could enable ascending feedback inhibition from the IGL to the OGL and eye. This work reveals fundamental differences between the visual systems of octopus and vertebrates, at both the neurochemical and microcircuit levels.

## Results

### Neurotransmitter usage in the eye and optic lobe

To investigate neuronal signaling in *Octopus vulgaris*, we focused on the newly-hatched paralarva, or hatchling, as a tractable model. A single *O. vulgaris* spawning gives rise to >100,000 progeny, which are around 2 mm long at hatching. While the hatchling brain is smaller than that of the adult, with only ∼200,000 neurons, it has the same lobular structure, including a prominent optic lobe with a laminar cortex^18,19^. Moreover, transcriptomic studies indicate that the paralarval brain shares most neural classes found in older animals ^9,10,20^. Paralarvae rely heavily on vision for their survival, and they exhibit robust visually guided behaviours including phototaxis, escape, colour change and prey capture^21^.

We first quantified the spatial (co-)expression of all neurotransmitters in the eye and optic lobe in detail. Based on previous data on neurotransmitter expression in the OL^9^, we used RNA fluorescence *in situ* hybridisation chain reaction (HCR) to probe the expression of six neurotransmitter marker genes in the photoreceptors and the three layers of the OL. We used the vesicular ACh transporter (vacht) as a marker for ACh, tyramine beta hydroxylase (tbh) for octopamine, the vesicular GABA transporter (vgat) for GABA, tryptophan hydroxylase (tph2) for serotonin, tyrosine hydroxylase (th) for dopamine, and the vesicular glutamate transporter (vglut) for glutamate, and quantified (co-)expression in the optic lobe.

We observed highly specific patterns of expression for each neurotransmitter in the paralarval visual system. In the eye, we observed positive signal only for glutamate (Fig. 1 a-f), indicating that the photoreceptors are glutamatergic, consistent with earlier results showing expression of glutamate receptors in the plexiform layer^12^. In the OL, we confirmed robust expression of four neurotransmitters, categorising them into five cell types: cholinergic (Chol: vacht^+^), octopaminergic (Octo: tbh^+^), exclusively glutamatergic (Glut: vglut^+^/th^−^), exclusively dopaminergic (Dop: th^+^vglut^−^), and dopaminergic/glutamatergic co-expressing (Dop/Glut: th^+^vglut^+^). Chol cells were absent from the OGL but represented a moderate number of cells in the IGL (20%) and the highest proportion of cells in the medulla (59%) (Fig. 1 g and i, Supplemental Fig. 1 a-c). Octo cells were relatively uncommon and were found only in the OGL (10%) and IGL (4%) (Fig. 1 h and i, Supplemental Fig. 1 d-f). Glut cells were found in all layers, representing 7% of cells in the OGL, 32% of cells in the IGL and 55% in the medulla (Fig. 1 j and m, Supplemental Fig. 2 a-c). Dop cells were the main cell type in the OGL (64%) and were also detected infrequently in the IGL (6%) and medulla (2%) (Fig. 1 k, m, Supplemental Fig. 2 d-f). Dop/Glut cells, which expressed both dopamine and glutamate, a cell type observed in transcriptomic studies from both hatchling and adult brains^9,10^, were the main cell type in the IGL (47%) and were less frequently seen in the OGL (11%) and medulla (4%) (Fig. 1 l and m, Supplemental Fig. 2 g-i). GABA and serotonin markers were detected only in a small number of medulla neurons, whose numbers we did not quantify (Supplemental Fig. 2 j, k). In summary, the OGL was composed primarily of Dop neurons along with other minor cell types, the IGL was a mixture of Glut, Chol and Dop/Glut cells, and the medulla was mainly a mixture of Chol and Glut cells, consistent with previous studies in *O. vulgaris* and *O. bimaculoides*^9,10^.

**Fig. 1.**
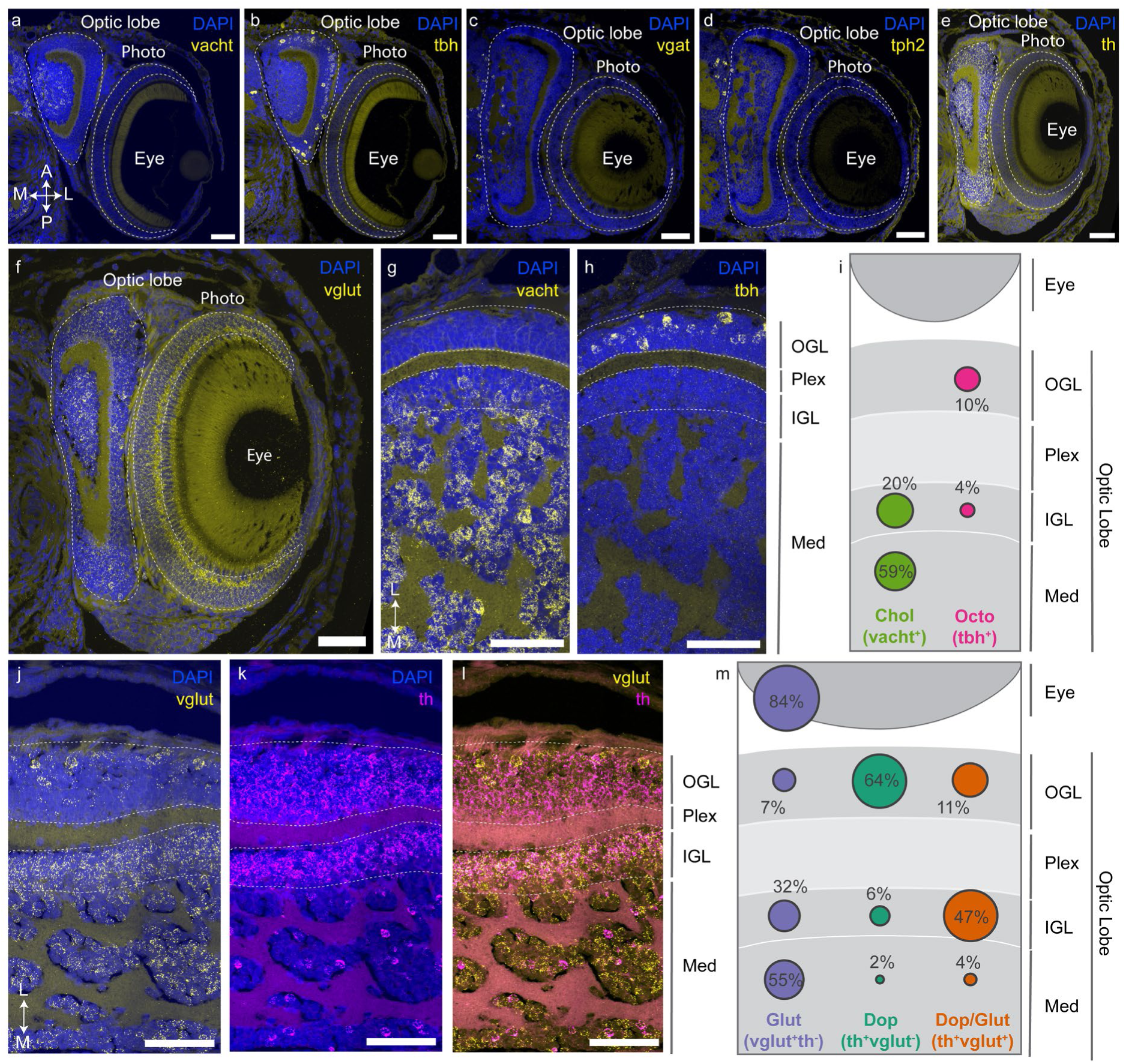
Neurotransmitter cell types in the eye and optic lobe of *O. vulgaris* paralarvae. HCR was performed on 1-dph *O. vulgaris* head transversal sections; nuclei were visualised in blue with DAPI. Sections with both the optic lobe and eye visible are from the ventral (towards the arms) part of the brain (see Supplemental Fig. 7 for more details). Representative HCR against vacht (cholinergic marker) (**A** and **G**), tbh (octopaminergic marker) (**B** and **H**), vgat (GABAergic marker) (**C**), tph2 (serotonergic marker) (**D**), th (dopaminergic marker) (**E** and **K**), and vglut (glutamatergic marker) (**F** and **J)** in yellow. Representative multiplex HCR against th (dopaminergic cell marker) in magenta and vglut (glutamatergic cell marker) in yellow (**L**). Summary of HCR analysis of Chol and Octo cell types in the eye and optic lobe (**I**). Summary of HCR analysis of Glut, Dop and Dop/Glut cell types in the eye and optic lobe (**M)** The size of the circle represents the proportion of cells in that layer belonging to that cell type. The head orientation is the same in A to F, G to M. Scale bar: 50 μm. vacht: vesicular acetylcholine transporter, tbh: tyramine beta hydroxylase, vgat: vesicular GABA transporter, tph2: tryptophan hydroxylase, th: tyrosine hydroxylase, vglut: vesicular glutamate transporter, OGL: outer granular layer, IGL: inner granular layer, Med: medulla, Plex: plexiform layer, Octo: octopaminergic cell type (tbh^+^), Chol: cholinergic cell type (vacht^+^), Dop: exclusively dopaminergic cell type (th^+^vglut^−^), Glut: exclusively glutamatergic cell type (vglut^+^/th^−^), Dop/Glut: exclusively dopaminergic and glutamatergic cell type (th^+^vglut^+^), A: anterior, P: posterior, M: medial, L: lateral.

### A dopamine-gated cation channel expressed in the optic lobe

We next sought to functionally characterise neurotransmitter receptors expressed in the optic lobe. We focused on cys-loop ligand gated ion channels (LGCs), a diverse family of ionotropic receptors that includes the GABA-A, glycine, 5-HT3, and the nicotinic ACh receptors of vertebrates as well as receptors with novel ligand specificity and ion selectivity found in some invertebrate species^14–17^. We identified putative *Octopus* cys-loop LGCs using comparative phylogenetics (Fig. 2 a) and used published single-cell RNA sequencing (scRNA-seq) data ^9^ to focus on those expressed in the optic lobe. We expressed 41 of these candidate *O. vulgaris* ‘cys-loop’ LGCs in *Xenopus* oocytes and used two-electrode voltage clamp (TEVC) recordings to measure currents evoked by application of 18 candidate ligands. Using this approach, we identified and characterised several candidate receptors activated by neurotransmitters expressed in the optic lobe (Supplemental file 1).

**Fig. 2.**
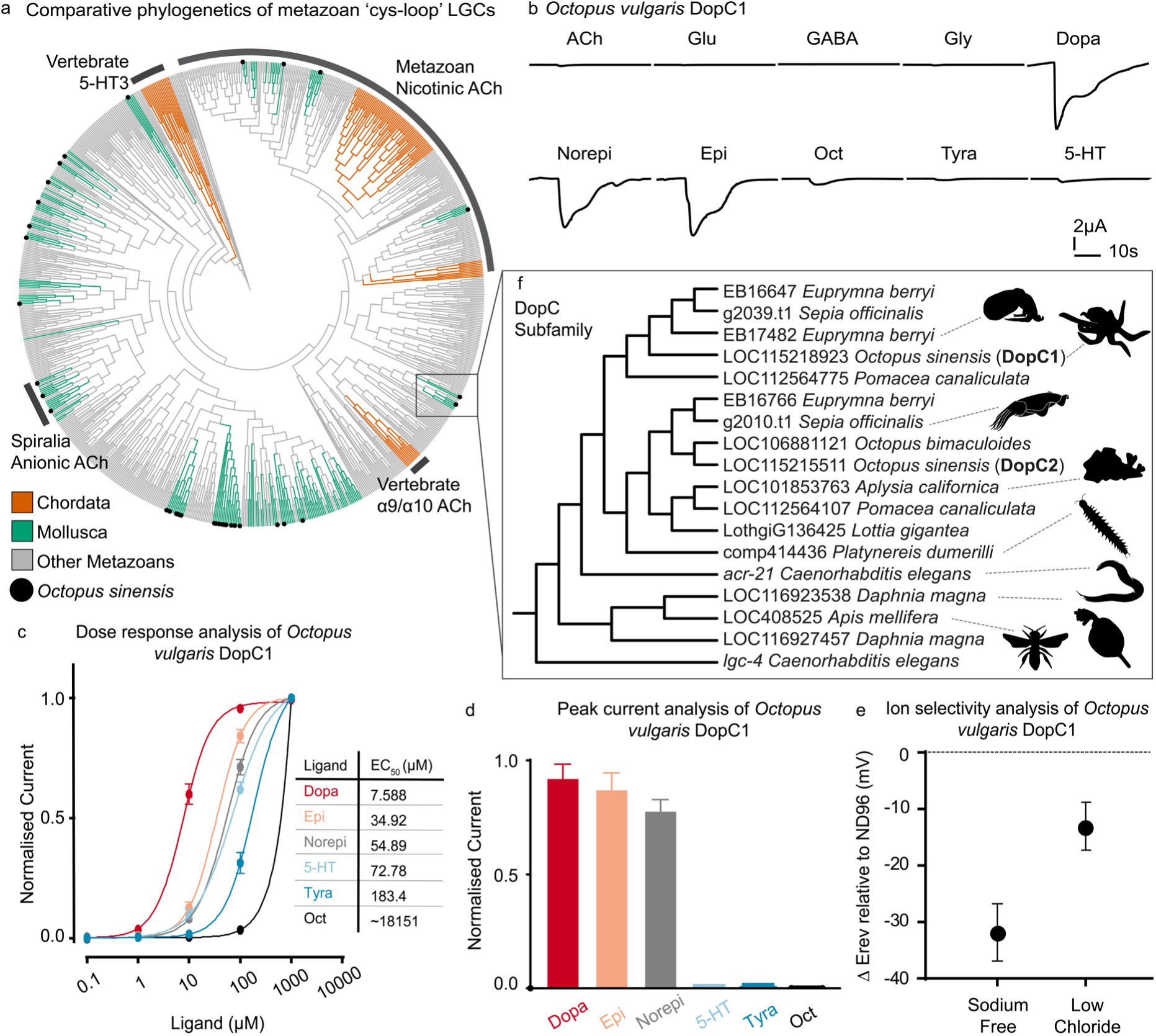
A dopamine-gated cation channel in *Octopus vulgaris* and other invertebrates. (**A**) Comparative phylogenetics of nicotinic-like cys-loop ligand gated ion channels from major metazoan phyla (cys-less and GABA-A like branches were collapsed). Chordata sequences are highlighted in orange and molluscan sequences are highlighted in green. *Octopus sinensis* sequences are highlighted with a black dot. A zoomed view of the DopC1 subfamily is in panel (**F**). (**B**). Representative TEVC traces from *Xenopus* oocytes expressing DopC1 from *O. vulgaris* during application of candidate ligands. ACh: acetylcholine, Glu: glutamate, Gly: glycine, Dopa: dopamine, Norepi: norepinephrine, Epi: epinephrine, Oct: octopamine, Tyra: tyramine, and 5-HT: serotonin. (**C**). Monoamine dose response analysis curves for *O. vulgaris* DopC1 expressed in *Xenopus* oocytes. The adjacent table includes the EC_50_ values for each ligand in µM. Mean and standard error are plotted for 3-7 separate oocytes per ligand. The current was normalised to the max current in each oocyte. (**D**). Peak current analysis (mean and standard deviation) of *O. vulgaris* DopC1 expressed in *Xenopus* oocytes. Each oocyte was exposed to 1 mM of the indicated monoamine. The current was normalised to the peak current in each oocyte. (**E**). Ion selectivity analysis of *O. vulgaris* DopC1 expressed in *Xenopus* oocytes during TEVC recordings Plotted is the difference in reversal potential (Erev, in mV) of the channel, measured during activation with 10µM (∼EC_50_) dopamine between ND96 and the sodium-free solution or the low-chloride solution. The mean and standard deviation are shown for 5 oocytes.

Among these putative receptors was a homomeric channel encoded by the LOC115218923 gene, which was activated by dopamine and other monoamines (Fig. 2 b and Supplemental Fig. 3 a). Expression analysis using scRNA-seq revealed that the LOC115218923 gene is expressed across the brain, in particular within cell-types of the optic lobe (Supplemental Fig. 3 b, d). In TEVC recordings of LOC115218923-expressing oocytes, we observed currents in response to dopamine (EC_50_: 7.6 μM), norepinephrine (55 μM), epinephrine (35 μM), serotonin (73 μM), tyramine (180 μM), and octopamine (18 mM)) (Fig. 2 c). This EC_50_ is consistent with a synaptic receptor, as extrasynaptic receptors localized further from the release sites often show an EC_50_ in the nM range. LOC115218923-expressing oocytes showed the highest mean peak current when exposed to dopamine, with norepinephrine and epinephrine (whose presence in the optic lobe has not been established) also showing large currents and serotonin, tyramine and octopamine giving negligible responses (Fig. 2 d).

Interestingly, a metabotropic dopamine receptor agonist, apomorphine ^22^, acted not as an agonist, but as an antagonist of dopamine-evoked responses (IC_50_: 7 µM, Supplemental Fig. 4 a and c). When we performed ion substitution experiments, comparing the shift in reversal potential (ΔErev) for a sodium-free or low-chloride solution compared to the standard buffer (ND96), we observed a large reversal potential shift in the sodium-free solution and little reversal shift in the low chloride-solution (Fig. 2 e). Thus, LOC115218923 appears to form a dopamine-activated cation channel, which we designated DopC1 (for ***<u>dop</u>****amine-gated **<u>c</u>**ation* channel).

Comparative phylogenetic analysis from diverse metazoan species indicated that DopC1 has homologues throughout bilateria. In vertebrates its closest homologues are the α9/α10 nicotinic receptors, cation channels that are activated by ACh but not dopamine (Fig. 2 a).

DopC1 was not activated by ACh; moreover, nicotine, a potent antagonist for mammalian α9/α10 receptors and agonist for other nicotinic channels, had no effect on DopC1-expressing oocytes as either an agonist or antagonist (Supplemental Fig. 4 b and c). We also found direct orthologs to DopC1 in many invertebrate species, including other cephalopods (*Euprymna berryi*, *Sepia officinalis*), molluscs (*Pomacea canaliculata, Aplysia californica,* and *Lottia gigantea*), an annelid (*Platynereis dumerilii*), a nematode (*Caenorhabditis elegans*), and some arthropods (*Apis mellifera* and *Daphnia magna,* but not *Drosophila melanogaster*). (Fig. 2 f). Expression of the DopC orthologs from *Aplysia* (LOC101853763), *Apis* (LOC408525) and *Daphnia* (LOC116927457 and LOC116923538) in *Xenopus* oocytes also produced homomeric channels activated by monoamines and not ACh (Fig. 3 a-d, Supplemental Fig. 5 a-d). For all these channels, we found that dopamine had the lowest EC_50_, though in some cases other monoamines such as norepinephrine, serotonin and tyramine also showed an EC_50_ in the µM range suggesting they might act as secondary ligands (Fig. 3 e-h). Peak current analysis on oocytes expressing these invertebrate DopC1 orthologs likewise indicated that dopamine, norepinephrine and epinephrine evoked the largest peak currents (Fig. 3 j). We also found a second DopC1-related sequence in *O. vulgaris*, but this gene (LOC115215511/DopC2), while expressed in the optic lobe (Supplemental Fig. 3 c and d), did not form a functional homomeric channel when expressed in *Xenopus* oocytes (Supplemental Fig. 5 e). Taken together, these results suggest that in addition to its well-established neuromodulatory action through metabotropic receptors, dopamine may act as a fast excitatory neurotransmitter through ionotropic receptors in octopus.

**Fig. 3.**
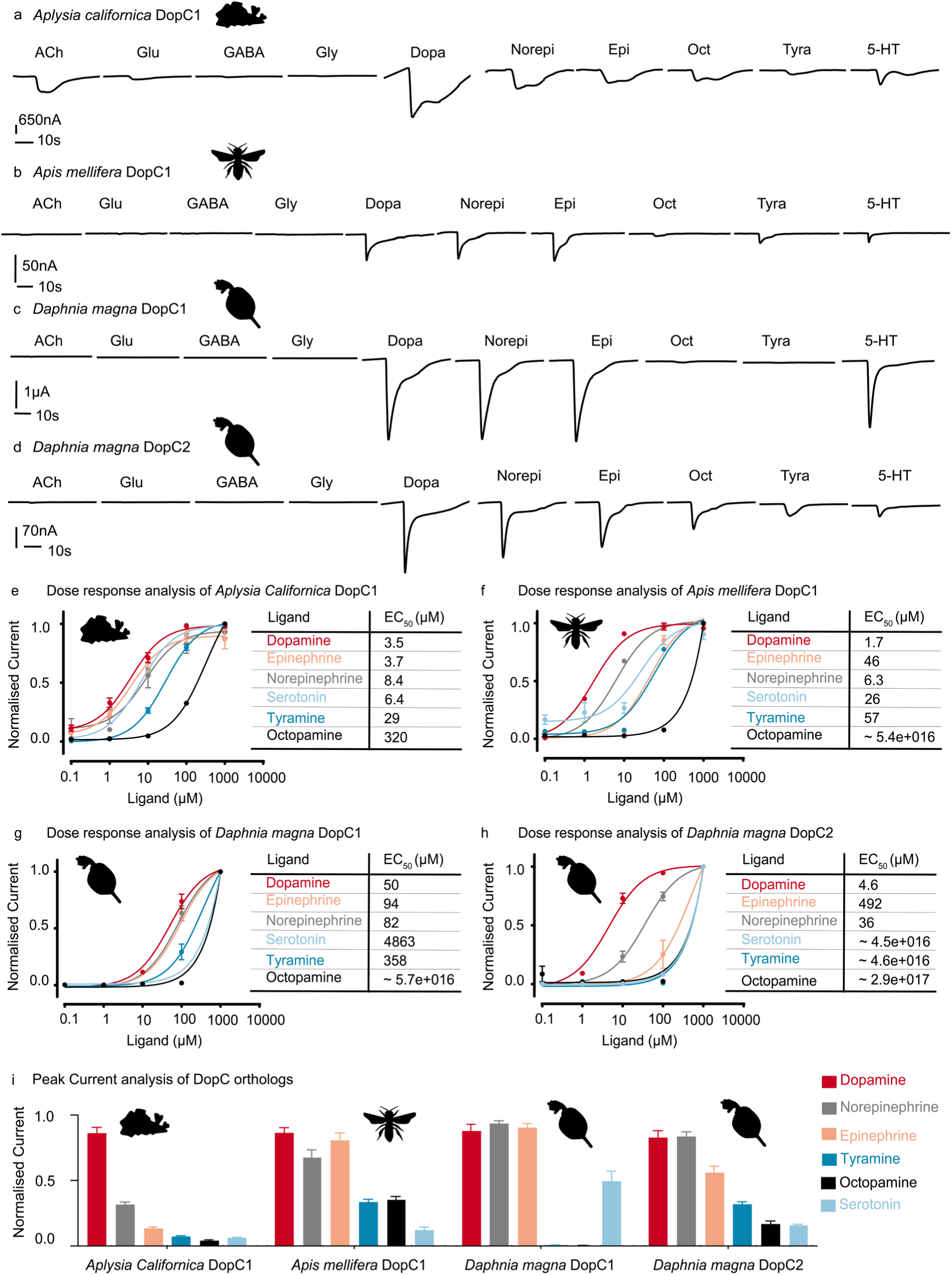
Functional characterisation of DopC orthologs from *Aplysia californica, Apis mellifera* and *Daphnia magna.* Representative TEVC traces from *Xenopus* oocytes expressing DopC1 from *Aplysia californica* (**A**), DopC1 from *Apis mellifera* (**B**), DopC1 from *Daphnia magna* (**C**), and DopC2 from *Daphnia magna* (**D**) during application of candidate ligands. Dose response analysis curves for the major monoamines during TEVC recordings of *Xenopus* oocytes expressing DopC1 from *Aplysia californica* (**E**), DopC1 from *Apis mellifera* (**F**), DopC1 from *Daphnia magna* (**G**), DopC2 from *Daphnia magna* (**H**). The adjacent table includes the EC_50_ values for each ligand in µM. Mean and standard error are plotted for 3-7 separate oocytes per ligand. The current was normalised to the max current in each oocyte. Peak current analysis for homomeric DopC orthologs from *Aplysia californica*, *Apis mellifera* and *Daphnia magna* expressed in *Xenopus* oocytes (**I**). During TEVC recordings each oocyte was exposed to 1mM dopamine, epinephrine, norepinephrine, serotonin, tyramine and octopamine. The mean and standard error of the mean are plotted per ligand, per receptor. The current has been normalised to the peak current in each oocyte. ACh: acetylcholine, Glu: glutamate, Gly: glycine, Dopa: dopamine, Norepi: norepinephrine, Epi: epinephrine, Oct: octopamine, Tyra: tyramine, 5-HT: serotonin, DopC: dopamine-gated cation channel.

### Dopaminergic excitation in the optic lobe

To explore the possible role of dopamine as a fast excitatory transmitter in the visual system, we examined the spatial expression pattern of DopC1 in the hatchling brain of *O. vulgaris,* with the goal of identifying cells whose activity might be stimulated by dopaminergic neurotransmission. We performed quantitative HCR using probes for DopC1, either alone (Fig. 4 a and b) or co-labelled with marker genes for the major optic lobe neurotransmitters (Fig. 4 e-h, Supplemental Fig. 6 a-e). In these experiments we detected no expression of DopC1 in the eye (Fig. 4 a), but observed widespread expression in the optic lobe (Fig. 4 a and b), principally in the IGL and medulla. Specifically, DopC1 was expressed in 21% of OGL neurons, 72% of IGL neurons, and 53% of medullar neurons (Fig. 4 b-d). In the OGL, DopC1 expression was disproportionately observed in the minor cell types (Glut, Octo, or Dop/Glut) and was rare in the predominant Dop cell type (Fig. 4 i, Supplemental Fig. 6 f). DopC1 showed the highest expression in the IGL, where it was expressed in a majority of each of the three major cell types in the IGL: 60% of Glut cells, 71% of Chol cells, and 92% of Dop/Glut cells, the most common class in this layer (Fig. 4 i, Supplemental Fig. 6 g).

**Fig. 4.**
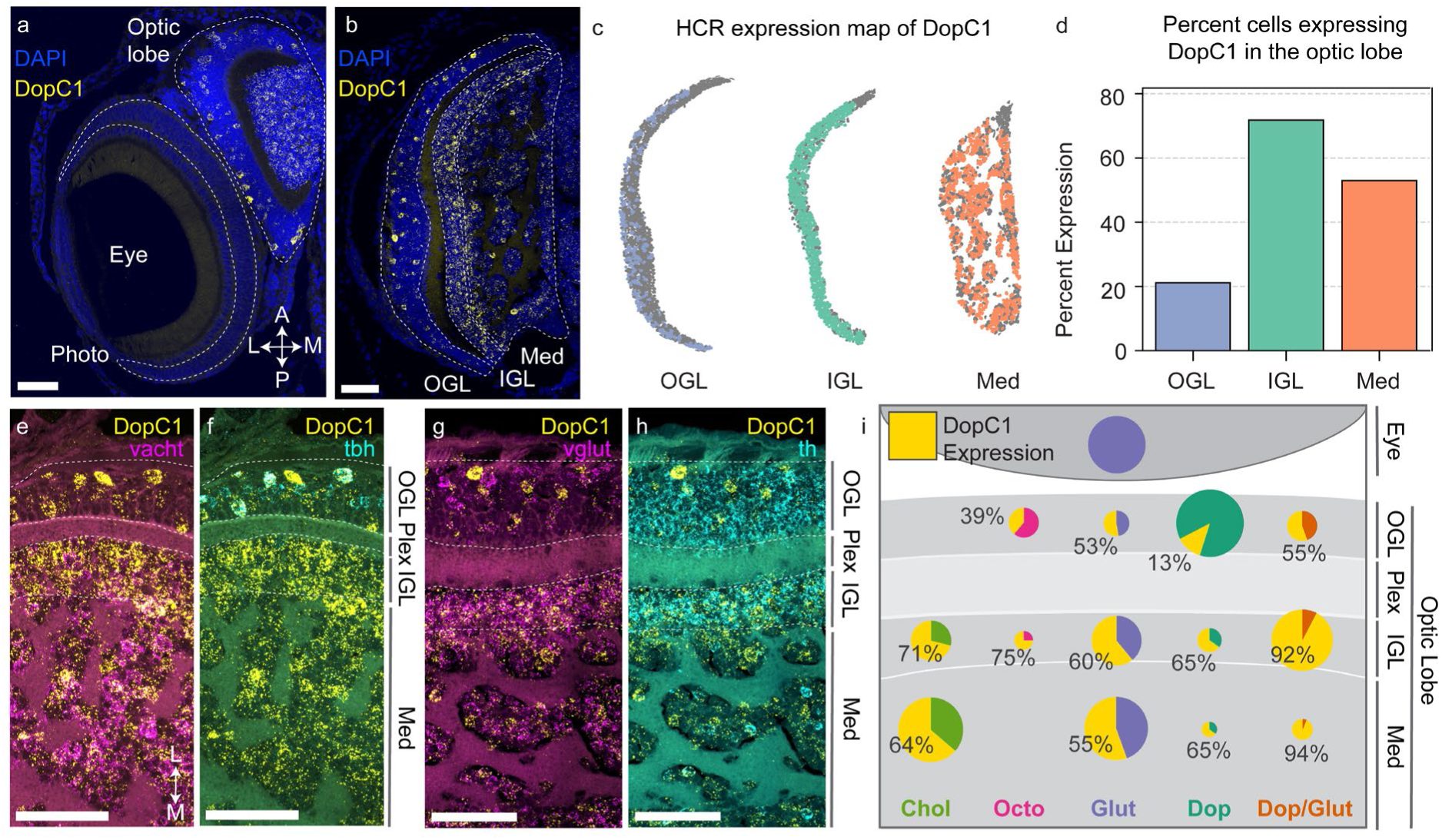
DopC1 expression in the eye and optic lobe of *O. vulgaris* 1dph paralarvae. (**A** and **B**) Representative hybridisation chain reaction (HCR) analysis of DopC1 expression. HCR was performed on 1-dph *O. vulgaris* head transversal sections; DopC1-expressing cells are in yellow; nuclei were visualised in blue with DAPI. (**C** and **D**) HCR expression maps (**C**) and barplot (**D**) showing the proportion and spatial distribution of cells expressing DopC1 in the OGL, IGL and Med of the OL, quantified from the HCR shown in B. (**E**-**H**) Representative multiplex HCR against DopC1 and indicated neurotransmitter markers. DopC1 is in yellow (all panels); vacht (cholinergic cell marker) in magenta (**E**), tbh (octopaminergic cell marker) in cyan (**F**), vglut (glutamatergic cell marker) in magenta (**G**) and th (dopaminergic cell marker) in cyan (**H**). The head orientation in A-C and E-I is the same. Scale bar: 50 μm. A: anterior, P: posterior, M: medial, L: lateral. (**I**) Summary of the expression of DopC1 in the major neurotransmitter cell types in the eye and OL. OGL: outer granular layer, IGL: inner granular layer, Med: medulla, Plex: plexiform layer, Octo: octopaminergic cell type (tbh^+^), Chol: cholinergic cell type (vacht^+^), Dop: exclusively dopaminergic cell type (th^+^vglut^−^), Glut: exclusively glutamatergic cell type (vglut^+^/th^−^), Dop/Glut: dopaminergic and glutamatergic cell type (th^+^vglut^+^). The size of the circle representatives the proportion of cells in that layer that belong to that cell type; the yellow sector represents the proportion of cells of that cell type that express DopC1. This quantification was performed on the multiplex HCRs shown in Supplemental Fig. 6.

DopC1 was also detected in a majority of cells of the major medullar cell types, namely Chol (64% DopC1 expressing) and Glut (55% DopC1 expressing). Other cell types with and without DopC1 represented a very low percent of cells in the medulla (1-4%; Fig. 4 i, Supplemental Fig. 6 h). DopC1 was also expressed throughout the central brain but at lower levels compared to the optic lobe (Supplemental Fig. 7). In summary, DopC1 showed relatively low expression in the OGL, but widespread expression in the IGL and medulla.

Based on this expression pattern, we hypothesised that dopamine might act as an excitatory transmitter to stimulate activity in deeper layers of the OL. To test this hypothesis, we performed calcium imaging on acute brain slices from 1-day *O. vulgaris* paralarvae, measuring individual cell responses to exogenous dopamine (Supplemental Fig. 8 a). We stimulated brain slices with 100 µM dopamine, followed by a high potassium solution (HighK^+^) as a control to identify actively-excitable cells, and characterized responses of individual cells to dopamine. These experiments indicated that dopamine has a predominantly excitatory role in the optic lobe (Fig. 5 a, b); excitatory responses to dopamine were seen in all layers of the optic lobe, while inhibitory dopaminergic responses were rare in all layers (≤1%). Notably, the proportion of cells showing excitatory dopaminergic responses, and the kinetics of these responses, varied across the different optic lobe layers. For example, a majority of cells in the IGL and the medulla showed positive responses to dopamine (83% and 68% respectively), while many fewer cells in the OGL (44%) showed dopaminergic excitation. Thus, excitatory dopamine responses were most common in the layers of the optic lobe with pervasive DopC1 expression, and rarest in the OGL where DopC1 was most sparse.

**Fig. 5.**
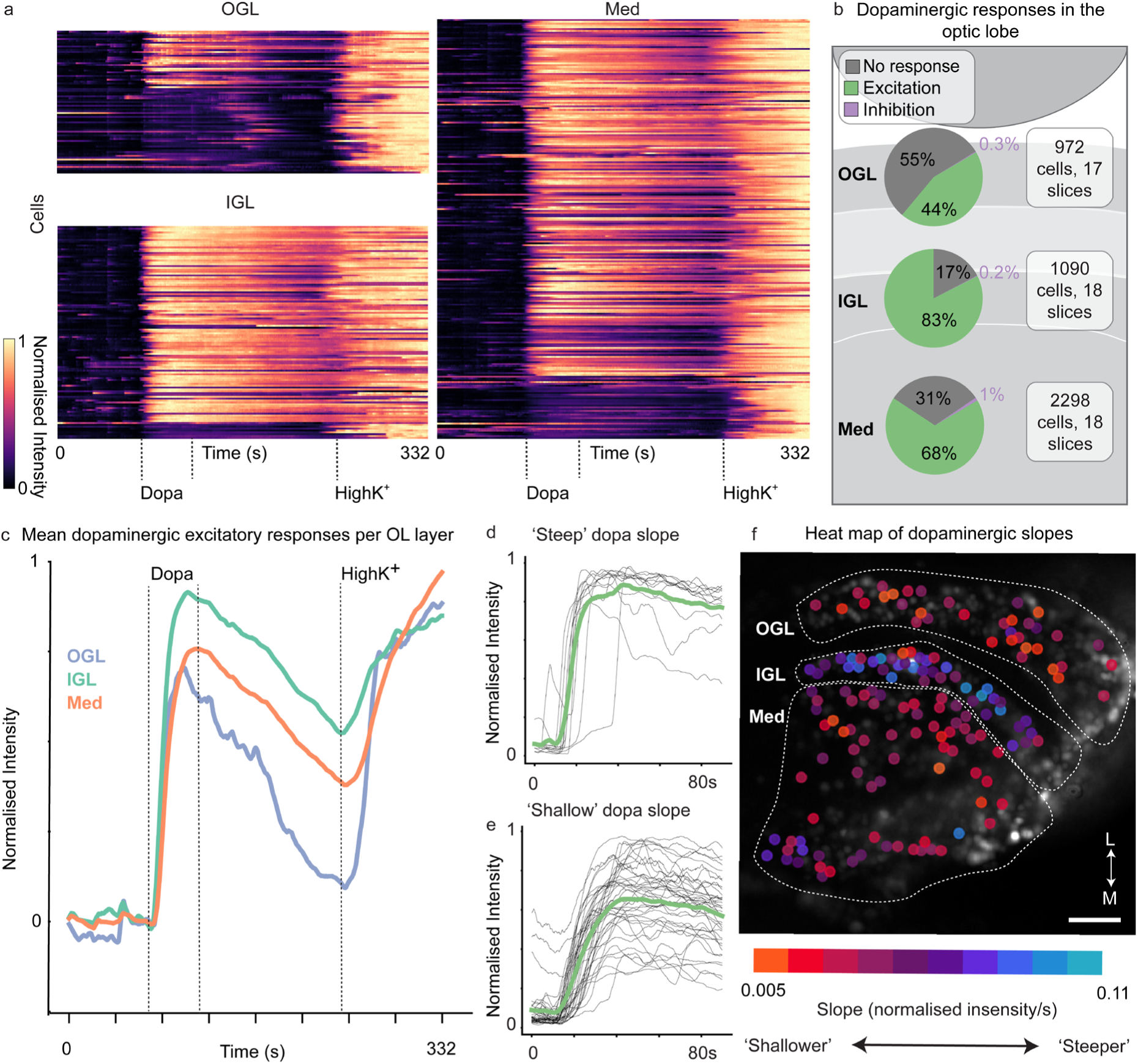
Dopamine excites neural activity in *O. vulgaris* paralarvae acute brain slices. (**A**). Calcium imaging was performed on 1-dph *O. vulgaris* acute brain slices during application of 100 µM dopamine (Dopa) and the HighK^+^ solution. Shown is a heat map of a representative slice during calcium imaging, with lines corresponding to individual cells in the indicated layer. The fluorescence intensity for each ROI was smoothed using a rolling average window over 10 seconds and normalised using min/max values per ROI. (**B**) Pie charts showing the proportion of cells which have dopaminergic excitation, dopaminergic inhibition or no response to dopamine, within the different layers of the optic lobe, summed across all brain slices (**C**). Mean traces from the representative slice shown in A per optic lobe layer (**D**). A subset of dopaminergic responses with ‘steep’ slopes ( >0.075 normalised intensity/s) were characterised as fast-rising responses (**E**). A subset of dopaminergic excitatory responses with “shallow” slopes (between 0.0125 and 0.0375 normalised intensity/s) were classes as slower responses. For responses in D and E, the mean trace is shown in green. (**F**). Maximum intensity projection of a representative brain slice with coloured dots indicating the slope values of the excitatory dopaminergic responses in the different layers of the optic lobe. ‘Steep’, fast-rising responses were most prevalent in the IGL, where DopC1 is most extensively expressed. Scale bar: 50 µm;, M: medial, L: lateral.

When we analysed the responses more closely, we also observed differences between the layers in the kinetics of dopamine response, reflected in the slope, relative amplitude, and duration. In particular, the IGL showed the highest dopamine response amplitudes (relative to HighK^+^ response) compared to the other layers. Likewise, the IGL, along with the medulla, showed longer lasting responses on average compared to the OGL (Fig. 5 c, Supplemental Fig. 8 b-e). Finally, the slope of dopaminergic responses also varied between cells, with all layers containing cells with ‘shallow’ slopes, indicating slow-increasing responses, as well as those with steeper slopes, indicating fast-increasing responses (Fig. 5 d and e). The cells of the IGL showed statistically steeper dopamine response slopes compared to the OGL and medulla (Games-Howell post-hoc test, p<0.0001), and a higher proportion of cells with steeply-rising slopes (Fig. 5 f, Supplemental Fig. 8 f). Thus, the rapid responses most consistent with direct ionotropic activation by dopamine were most prevalent in the IGL, the layer where DopC1 expression is most prevalent (Fig. 4 d). These results are consistent with dopamine acting through DopC1 as a fast excitatory neurotransmitter in the IGL, and perhaps also in the medulla.

### An acetylcholine-gated anion channel expressed in dopaminergic neurons

To gain insight into how dopaminergic neurotransmission in the optic lobe is regulated, we next focused on an LGC with specific expression in dopaminergic neurons. Analysis of scRNA-seq data^9^ revealed that a gene related to nicotinic ACh receptors, LOC115214201, showed high and specific expression in dopaminergic clusters (Supplemental Fig. 9 a and b). To investigate the functional properties of this channel, we expressed it in *Xenopus* oocytes and performed TEVC recordings (Fig. 6 a, and Supplemental Fig. 10 c). We observed that oocytes expressing the LOC115214201 gene, hereafter referred to as AChRB1, showed currents in response to both ACh and choline, though the EC_50_ of 1.9μM for ACh was much lower than that for choline (98 μM) (Fig. 6 b). This indicates that ACh likely acts as the primary endogenous ligand, and the micromolar EC_50_ for ACh is consistent with AChRB1 acting as a synaptic receptor. To determine the ion selectivity of AChRB1, we performed ion substitution experiments, as described previously for DopC1. For these experiments we used the AChRB1 ortholog from *Octopus bimaculoides,* as it expressed more robustly in oocytes compared to the *O. vulgaris* gene while sharing the same ligand specificity properties (Supplemental Fig. 10 d and e). We found that the AChRB1 channel showed a large and significant shift in reversal potential in the low-chloride solution; in contrast, replacement of the normal buffer with a sodium-free solution caused no significant shift (Fig. 6 c). Thus, octopus AChRB1 appears to function as an anionic channel capable of mediating inhibitory cholinergic neurotransmission.

**Fig. 6.**
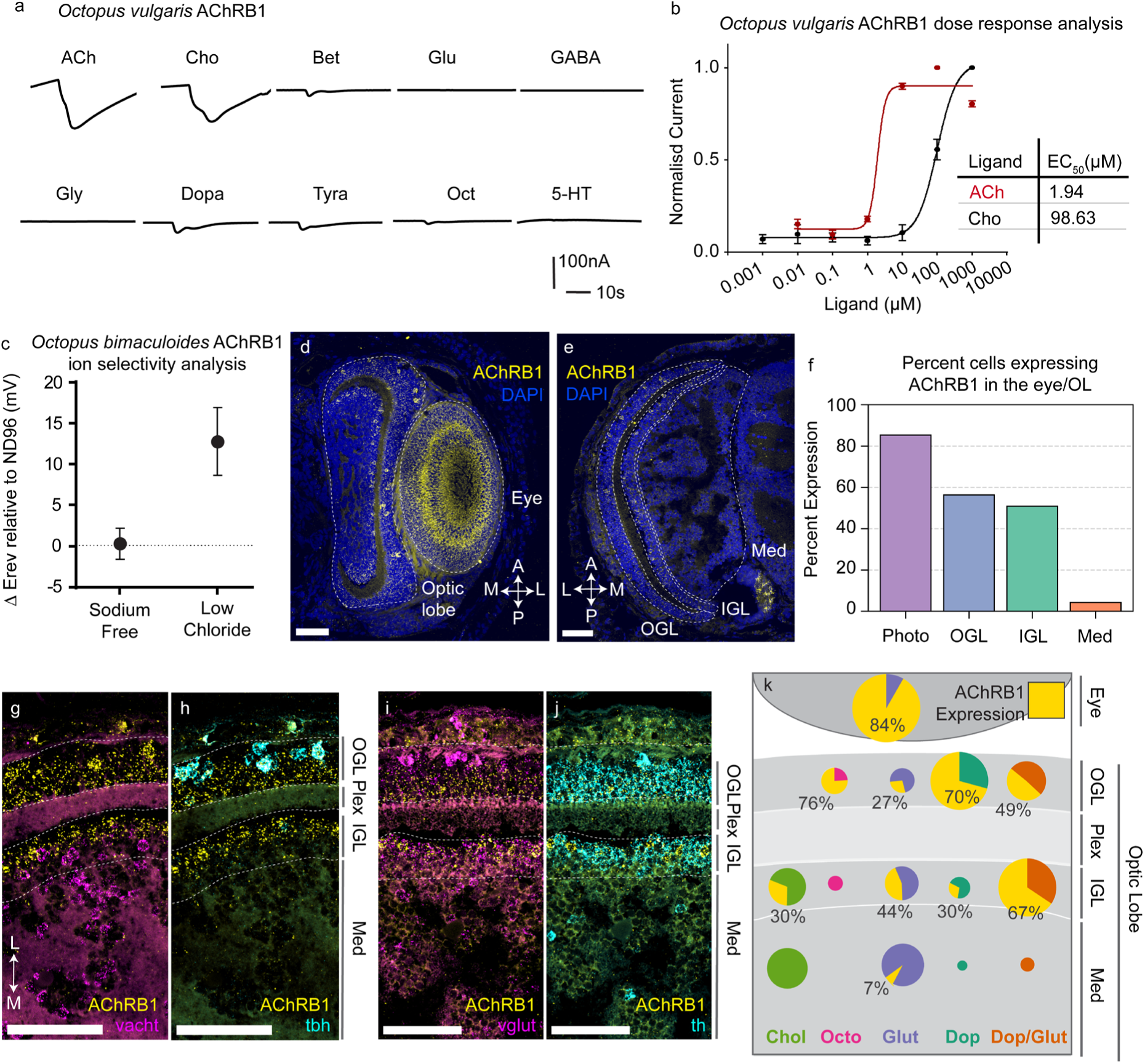
AChRB1, an ACh-gated chloride channel expressed in the *O. vulgaris* visual system. (**A**) Representative traces from *Xenopus* oocytes expressing *O. vulgaris* AChRB1 upon application of candidate ligands. Ligand abbreviations are as in Fig 2. (**B**) Dose response curves for ACh and choline for *O. vulgaris* AChRB1 expressed in *Xenopus* oocytes. EC_50_ values for each ligand are shown in the table. Mean and standard error are plotted for 5-6 separate oocytes per ligand; the current was normalised to the max current in each oocyte. (**C**) Ion selectivity analysis of *O. bimaculoides* AChRB1 expressed in *Xenopus* oocytes. Plotted is the difference in reversal potential (Erev, in mV) of the channel upon activation with 1µM ACh (∼EC_50_) between ND96 and either sodium-free solution or low-chloride solution. The mean and standard deviation are shown for 7 oocytes. (**D** and **E**) Representative HCR analysis of AChRB1 expression. All HCRs were performed on 1-dph *O. vulgaris* head transversal sections; AChRB1-expressing cells are in yellow, nuclei were visualised in blue with DAPI. (**F**) Barplot showing the proportion of cells expressing AChRB1 in the eye and OL, quantified from the HCR shown in D and E (**G-J**) Representative multiplex HCR against AChRB1 and indicated neurotransmitter markers; AChRB1 is in yellow (all panels); vacht (cholinergic cell marker) in magenta (**G**), tbh (octopaminergic cell marker) in cyan (**H**), vglut (glutamatergic cell marker) in magenta (**I**) and th (dopaminergic cell marker) in cyan (**J**). The orientation in all panels is the same; scale bar: 50 μm. (**K**) Summary of AChRB1 expression of AChRB1 in the eye and OL. Cell type and layer abbreviations are as in Fig 1. The size of the circle represents the proportion of cells in that layer that belong to that cell type; the yellow sector represents the proportion of cells of that neurotransmitter type that also express AChRB1. This quantification was performed on the multiplex HCRs shown in Supplemental Fig. 11.

Our comparative phylogenetic analysis of cys-loop LGCs from metazoans placed this gene within a group of anionic ACh-gated channels previously functionally characterised in the mollusc *Lymnaea stagnalis*^23^ (AChRB), and the platyhelminth *Schistosoma mansoni*^24^ (Fig. 2 a). Our phylogenetic analysis also found direct orthologs in other molluscs (*Aplysia californica, Pomacea canaliculata, Lottia gigantea*), platyhelminths (*Schmidtea mediterranea*) and annelids (*Platynereis dumerilii, Capitella teleta*). Despite being related to cationic nicotinic ACh-gated channels, all genes within this group contain a PAK (proline-alanine-lysine) motif in the pore lining region similar to that found in anion-permeable GABA-A and glycine receptors (Supplemental Fig. 9 c) that has been linked to chloride selectivity^25^. Thus, the AChRB family may form ACh-gated anion channels in many Lophotrochozoan phyla. Our phylogenetic analysis identified two other members of this family (LOC115229990/AChRB2 and LOC115212977/ AChRB3) in *O. vulgaris*, however, these did not show currents when expressed as homomeric channels in oocytes (Supplemental Fig. 10 a and b).

### Acetylcholine acts as an inhibitory transmitter in the optic lobe

To explore the effect of AChRB1 on octopus visual circuits, we first investigated its expression pattern in the brain of *O. vulgaris* paralarva through quantitative HCR. We observed widespread expression of the AChRB1 gene in the eye (85% of photoreceptor cells), in the OGL and IGL (56% and 51% respectively), and in a low proportion of medulla cells (4%) (Fig. 6 d-f). To investigate the expression pattern of AChRB1 in the optic lobe in more detail, we performed multiplex HCR of AChRB1 in combination with marker genes for visual system neurotransmitters (Fig. 6 g-j, Supplemental Fig. 11 a-e). Within the OGL, AChRB1 was expressed in most of the Dop neurons (70%), the most common cell type in the layer. It was also expressed a majority of the Octo cells (76%), as well as in around half of all Dop/Glut OGL cells (49%; Fig. 6 k, Supplemental Fig. 11 f). In the IGL, AChRB1 was expressed in most Dop/Glut cells (67%) the most common cell type in the layer, in a smaller fraction of Chol (30%) and Glut cells (44%), and not at all in Octo cells (Fig. 6 k, Supplemental Fig. 11 g). AChRB1 was expressed in very few medulla cells (7% of Glut cells) and some central brain cells (Fig. 6 k, Supplemental Fig. 11 h, Supplemental Fig. 12). To summarise, we observed AChRB1 expression most prominently in three cell types: Glut photoreceptors in the eye, Dop amacrine cells in the OGL, and Dop/Glut cells in the IGL.

Since all three of these cell types are potential recipients of synaptic input from cholinergic neurons in the IGL and/or the medulla^26^, and since scRNA-seq data indicated that cationic nicotinic ACh-gated channels are not highly expressed in dopaminergic clusters (Supplemental Fig. 13 a), we hypothesised that ACh might act through AChRB1 as an inhibitory neurotransmitter in the octopus visual system. To test this hypothesis, we performed calcium imaging on acute brain slices from *O. vulgaris* paralarvae following acute application of ACh, using the same protocol used for dopamine. We indeed observed mostly inhibitory effects of ACh on optic lobe calcium activity (Fig. 7 a-e, Supplemental Fig. 14).

**Fig. 7.**
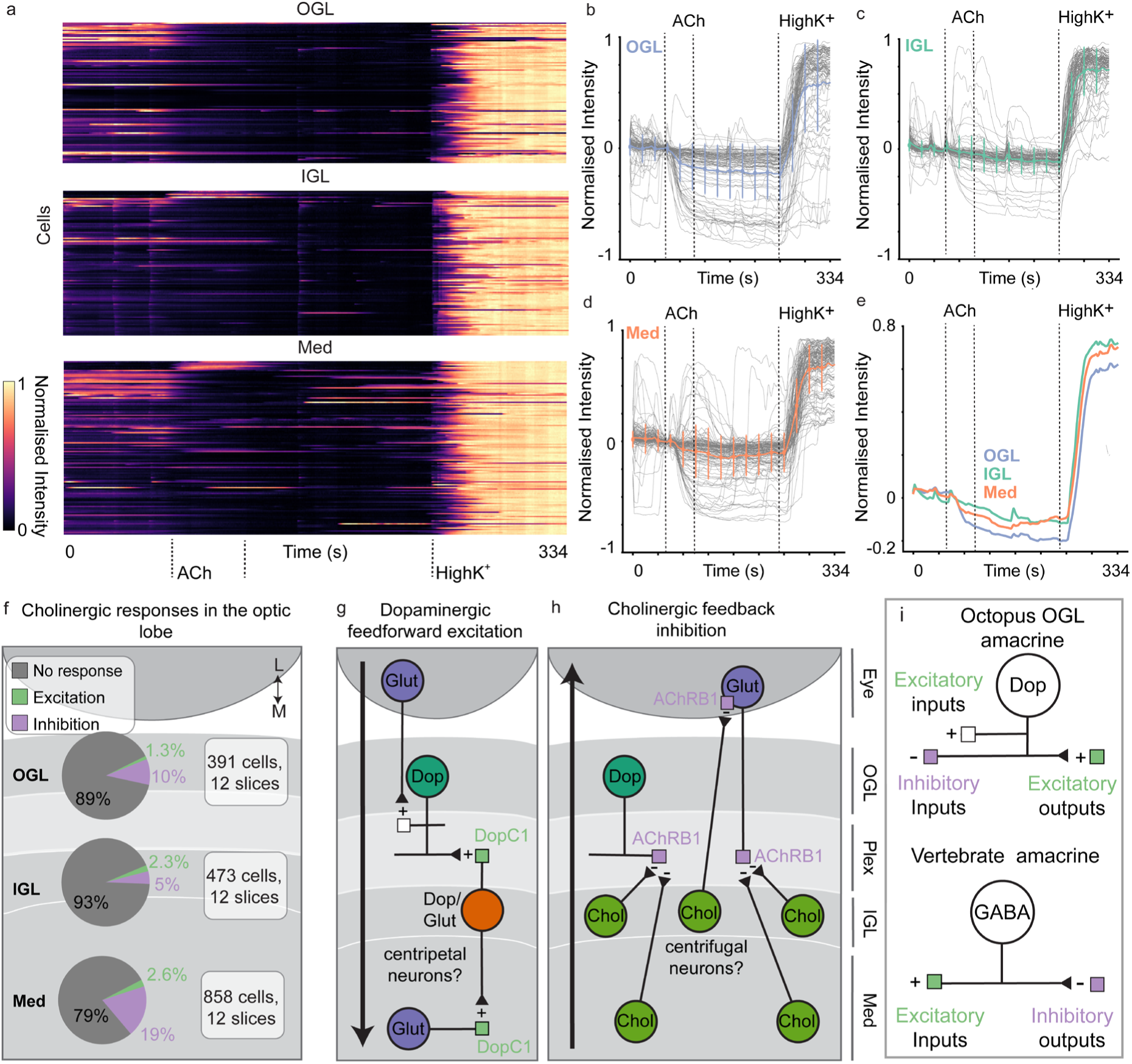
Acetylcholine inhibits neural activity in *O. vulgaris* paralarval acute brain slices. (**A**) Calcium imaging was performed on 1-dph *O. vulgaris* acute brain slices during application of 100 µM ACh (ACh) and the HighK^+^ solution. Shown are heat maps of a representative brain slice, with lines corresponding to individual cells in the indicated layer. The fluorescence intensity for each ROI was smoothed using a rolling average window over 10 second periods and normalised using min/max values per ROI. **(B-E)** Normalised intensity over time for all ROIs of the representative slice in panel A in the OGL (**B**), IGL (**C**) and Med (**D**). Mean trace and error bars are highlighted in bold. Panel **E** shows the mean traces from B-D overlayed on each other (**F**). Pie charts showing the proportion of cells which showed excitation, inhibition or no response to ACh, within the different layers of the OL, summed across all brain slices (**G-I**). Circuit hypotheses. Cell type and layer abbreviations are as in Fig 1. (**G**) Dopamine is hypothesises do carry out feedforward excitation between the OGL and IGL, and between IGL and Med. (**H**) Acetylcholine is hypothesised to carry out feedback inhibition between the IGL and/or Med and the OGL and/or photoreceptors. (**I**). Differences in input and output valences between octopus OGL dopaminergic amacrine cells and vertebrate retinal GABAergic amacrine cells.

ACh-mediated inhibition was observed in all layers of the OL: 10% of OGL cells, 5% of IGL cells, and 19% of medulla cells (Fig. 7 f); in contrast, excitatory responses to ACh were observed in less than 2.6% of all cells in each layer while other recorded cells had no detectable response (Fig. 7 f). The proportion of cells experiencing inhibition by ACh is likely underestimated in our experiments due to the absence of baseline activity which largely masks inhibition in our calcium imaging preparation. However, the fact that ACh elicited primarily inhibitory, rather than excitatory, responses in the optic lobe is consistent with AChRB1 mediating inhibitory neurotransmission in visual circuits.

## Discussion

In this study, we have begun to map the microcircuit architecture and function of the visual system of the *Octopus vulgaris* paralarva, with the goal of linking neurotransmitters, their receptors, and their physiological effects. In particular, we have defined the physiological properties of receptors for key optic lobe neurotransmitters, dopamine and ACh, and localised receptor expression relative to their neurotransmitter ligands within the eye and optic lobe. Through imaging of dopamine and ACh responses in different layers of the optic lobe, this work has provided new insights into the molecular and circuit mechanisms underlying visual processing in octopus.

These experiments revealed that dopamine, unexpectedly, functions as a major excitatory neurotransmitter in the optic lobe. Calcium imaging of octopus brain slices following application of dopamine showed acute activation in all optic lobe layers, most prominently in the IGL. This activation correlates with the expression of DopC1, a newly-characterised dopamine-gated cation channel found in nearly all IGL neurons, as well as some OGL neurons and about half of medullar neurons. Not only do the vast majority of IGL neurons (>80%) show increased activity in response to dopamine, the neurons in the IGL also showed consistently faster-increasing responses, suggesting direct activation via ionotropic DopC1 receptors. Most neurons in the OGL are dopaminergic, with axons juxtaposed with the dendrites of DopC1-expressing IGL neurons in the plexiform layer. This strongly suggests that signaling between neurons in the OGL and higher-order neurons in the IGL involves dopaminergic excitatory transmission, mediated by DopC1.

The DopC receptor identified in this study defines a new class of monoamine-gated cation channel whose members likely play important roles in excitatory neurotransmission in invertebrates. Excitatory currents activated by dopamine have been described previously in a gastropod mollusc, *Helisoma trivolvis*^27^, but the molecular identity of the receptors mediating these currents was not known. Since the DopC1 homologue from *Aplysia* is also monoamine-activated, we hypothesise that a DopC-related channel might also be responsible for dopaminergic excitatory transmission in gastropods. Although dopamine-gated channels have been previously identified in *C. elegans*, these are anionic channels that belong to a GABAA-related LGC subfamily that is specific to nematodes^15,16,28^. In contrast, cationic DopC1 receptors are homologous to vertebrate α9/10 nicotinic receptors and appear to form monoamine-gated channels in phyla ranging from lophotrochozoans such as molluscs and annelids to ecdysozoans such as arthropods and nematodes. Our results therefore suggest that in octopus, as well in many other invertebrates, dopamine not only acts as a slow neuromodulator through G-protein coupled receptors, but also as a fast excitatory neurotransmitter through DopC-related channels.

Our observation that dopamine is largely excitatory in the *O. vulgaris* optic lobe was surprising given a previous report^13^ that cuttlefish optic lobe neurons showed a decrease in spontaneous activity in a small sample of optic lobe neurons following dopamine application. While the authors make no claims about directly evoked dopaminergic currents, we were unable to confirm or refute these observations in our calcium imaging preparation due to its inability to resolve spontaneous (subthreshold) events. Alternatively, since octopods and decapods are separated by 250 million years of evolution, it is possible that dopamine may play different roles in the cuttlefish and octopus optic lobes. Both *Euprymna berryi* (squid) and *Sepia officinalis* (cuttlefish) have DopC1 homologs^29,30^; thus, it will be interesting to investigate how dopamine signaling functions in the visual circuits of these more distantly-related cephalopod species.

In the context of the optic lobe circuitry, dopaminergic transmission through DopC1 appears to play a key role in descending excitatory pathways that propagate visual information from first-order to higher-order neurons. Previous work has shown that cephalopod photoreceptors (in contrast to those of vertebrates) are excited by light^7,31^, have large presynaptic output terminals in the superficial plexiform layer of the optic lobe that contact the dendrites of OGL neurons^26^. While these photoreceptors were previously thought to be cholinergic^11,31^, recent scRNA-seq data from squid^29^ along with our HCR results from *Octopus vulgaris* indicate that they release the excitatory neurotransmitter glutamate. Since most OGL neurons are dopaminergic, we infer that these neurons relay and/or filter depolarizing signals from the photoreceptors to the dendrites of DopC1-expressing IGL neurons found deeper in the plexiform layer. Dopaminergic transmission may thus facilitate the retinotopic ON pathways recently described in *O. bimaculoides* during light stimulation^7^. Since some IGL neurons may also receive direct, glutamatergic excitatory input from the photoreceptors as well as feed-forward, dopaminergic excitation from OGL neurons, dopaminergic signaling may serve to amplify the signal between the OGL and IGL, as was observed in *O. bimaculoides* calcium imaging experiments^7^.

Dopaminergic signaling may also be important to propagate excitatory signaling between the IGL and the medulla. In particular, the DopC1-expressing Dop/Glut IGL neurons produce two excitatory neurotransmitters, glutamate^12^ as well as dopamine. Many neurons in the medulla express DopC1, and many also undergo activation by dopamine, in some cases with rapid kinetics (Fig. 5 f). Since very few neurons in the medulla synthesize dopamine, the dopaminergic inputs to the medulla are therefore likely to come primarily from Dop/Glut neurons in the IGL. Thus, the Dop/Glut neurons may correspond to the centripetal neurons described by Young, serving as descending excitatory relays of visual information from second-order neurons of IGL to third-order neurons of the medulla (Fig. 7 g).

Our results also define a role for ACh as an inhibitory neurotransmitter in the octopus optic lobe. Although inhibition is intrinsically difficult to detect using calcium imaging, we observed clear evidence of cholinergic inhibition in all layers of the optic lobe; in contrast, we observed little evidence for cholinergic excitation. This cholinergic inhibition correlates, particularly in the OGL and IGL, with the expression of an ACh-gated anion channel, AChRB1. AChRB1 is a nicotinic receptor whose pore domain is selective for anions rather than cations and is homologous to previously described ACh-gated anion channels in the gastropod *Lymnaea* and the platyhelminth *Schistosoma*^23,24^. The expression of AChRB1 in the photoreceptors and neurons of the optic lobe cortex correlates with previous reports that ACh caused inhibition in the plexiform layer of *O. vulgaris* adult optic lobe slices, while nicotinic receptor agonists such as α-bungarotoxin and atropine had no effect^32^. We note that we also identified classical nicotinic cationic ACh-gated channels in *O. vulgaris*, though they are not highly expressed in the optic lobe. Interestingly, ACh-gated anion channels have also been found in nematodes, but these are unrelated to AChRB1 and more closely related to GABA-A-like receptors^17^; indeed, AChRB-like channels, unlike DopC-like channels, have been found only in lophotrochozoan nervous systems.

In contrast to dopamine and glutamate, which appear to function primarily in descending, feed-forward pathways in the optic lobe, acetylcholine appears to act in ascending, feedback pathways. For example, we observed widespread expression of AChRB1 in the OGL, particularly in Dop and Dop/Glut cells; thus, ACh likely inhibits neurons releasing the two major excitatory neurotransmitters in the OGL, dopamine and glutamate. Neither the OGL nor the retina contains cholinergic neurons; thus, the cholinergic input neurons that act on AChRB1-expressing neurons in the OGL must be IGL or medullar neurons that project processes into the plexiform layer. In addition, we observed AChRB1 expression in photoreceptors, and previous reports indicate that ACh indeed exerts inhibitory effects on photoreceptor terminals in the plexiform layer. Thus, cholinergic neurons from the IGL appear to inhibit both photoreceptor terminals and dopaminergic OGL neurons in the plexiform layer. In addition, if cholinergic neurons correspond to the centrifugal cells reported by Young ^26^, they might also inhibit photoreceptors in the retina itself, or provide lateral inhibition to neighbouring IGL cells that express AChRB1(Fig. 7 h).

Since cholinergic inhibition was also seen in the medulla, it is possible that acetylcholine may function in descending inhibitory pathways as well. In *Octopus bimaculoides*, it has been shown that OFF retinotopic pathways first appear in the medulla^7^; thus cholinergic inhibition might carry out the required sign inversion between the IGL and medullar layers. It should be noted however that AChRB1 itself does not show detectable expression in the medulla; thus, the cholinergic inhibition in medullar neurons may be mediated by a different, yet-unidentified inhibitory ACh receptor. Alternatively, AChRB1 might act indirectly to inhibit medullar neurons by inhibiting the output terminals of excitatory centripetal neurons projecting from the IGL, similar to its inferred role at photoreceptor terminals in the plexiform layer.

The octopus visual system, like those of other cephalopods, shares important common features with that of vertebrates; indeed, the camera-like eye and laminar optic lobe cortex (OGL and IGL, sometimes called the “deep retina”) are often cited as examples of convergent evolution. However, at the microcircuit level our work suggests important differences may exist between the wiring of the octopus and mammalian visual pathways. In particular, the expression of inhibitory AChRB channels in photoreceptors and lower-order dopaminergic neurons implies the existence of ascending, negative feedback pathways in the first steps of octopus visual processing, from the inner granular layer and/or medulla to the OGL and the eye. This is in marked contrast to the mammalian retina, where no ascending pathways are found in the first three layers of the visual circuitry. Our results also imply an inverted polarity for lateral signaling in at least some neurons involved in early visual processing. In particular, dopaminergic OGL amacrine cells direct excitatory outputs (to DopC1-expressing targets) and receive inhibitory inputs (from cholinergic neurons via AChRB1) in the plexiform layer. In contrast, laterally-projecting horizontal and amacrine cells of the vertebrate retina receive excitatory glutamatergic inputs and make inhibitory GABAergic outputs in the inner and outer plexiform layers^8^ (Fig. 7 i). Thus, the visual circuitry of *Octopus* may contain novel circuit motifs, such as cholinergic-dopaminergic inhibitory feedback loops and reverse-polarity amacrine cells, that differ from those found in the mammalian retina.

Coleoid cephalopods have become of great interest to neuroscientists due to their independent evolution of intelligence and innovative behaviours. In addition to their well-established differences in systems-level organisation, our work here has provided functional and evidence that cephalopod and vertebrate visual systems are also profoundly different at the neurochemical and microcircuit levels. As a result of evolutionary diversification of the ligand and ion selectivity of ionotropic receptors, the octopus uses dopamine as a fast excitatory neurotransmitter, and ACh as an inhibitory transmitter, in the optic lobe. Likewise, the circuitry that carries out the earliest stages of visual processing in octopus makes use of microcircuit motifs that are not found in the vertebrate retina, suggesting they may use innovative circuit mechanisms for processes such as object identification and motion detection. Going forward, it will be interesting to identify computational principles that are shared between the visual circuits of cephalopods and vertebrates, as well as those unique to the complex, “alien” brain of the octopus.

## Methods

### Data and Code availability

All custom code developed for this study has been deposited to GitHub (see below). The phylogenetic analysis data is available in supplemental file 2-4. The HCR probe sequences are available in supplemental file 5. All images used for quantitative HCR are shown in main and supplemental figures. All other data are available from the authors upon request.

### Octopus vulgaris paralarvae

Adult male and female wild caught *O. vulgaris* were maintained and mated in the Oceanographic Center of the Canary Islands Instituto Español de Oceanografía (IEO-CSIC, Tenerife, Spain) according to Almansa et al.^33^ and upon spawning embryos were shipped to Belgium. Embryos were then incubated until hatching in a closed system in the Laboratory of Developmental Neurobiology (KU Leuven), Belgium (ref). Animals used in this study were from spawnings that occurred between September 2021 and November 2024. All procedures involving adults and hatchlings were approved by the ethical board and competent authority on animal experimentation from CSIC (permit CEIBA 1377-2023 and 1610-2024) and KU Leuven (permit P080/2021), in compliance with Directive 2010/63/EU.

### Octopus genomes

At the conception of this project a high-quality genome/transcriptome did not exist for *Octopus vulgaris* and thus the genome/transcriptome of *Octopus sinensis* (*ASM634580v1*) was used for designing primers for PCR and probes for HCR (more details below). *Octopus sinensis* is the most closely related octopus species to *Octopus vulgaris* and it was reported that there is only a 3.21% difference between the mitogenomes ^34^. In addition, the *O. vulgaris* scRNA-seq ^9^ was performed on *Octopus vulgaris* 1-dph paralarval brains but was mapped to the *Octopus sinensis* genome. At later stages of the project a high-quality genome/transcriptome was published for *Octopus vulgaris* (*PRJEB61268,* ^34^ and was incorporated into the phylogenetic analysis.

### Comparative phylogenetics of metazoan LGCs

#### Identifying LGCs in Octopus and diverse metazoan phyla

Cys-loop LGC protein sequences were identified using the PFAM domain PF02931, and the following publicly available sequence datasets were selected to include species representatives from all the major phyla. *Homo sapiens* (GRCh38.p14), *Danio rerio* (GRCz11), and *Ciona intestinalis* (GCA_000224145.1), were included as Chordate representatives. *Saccoglossus kowalewskii* (PRJNA42857) was the hemichordate representative. *Strongylocentrotus purpuratus* (Spur_5.0) was the Echinoderm representative. *Octopus sinensis (ASM634580v1)*, *Octopus bimaculoides (ASM119413v2)*, *Octopus vulgaris (PRJEB61268)*, *Euprymna berryi* (PRJEB52690), *Sepia Officinalis* (PRJNA1091451), *Aplysia californica (AplCal3.0)*, *Lottia gigantea (Lotgi1)* and *Pomacea canaliculate* (ASM307304v1) represented the Mollusca. Three octopus genomes were used to ensure we didn’t miss any LGCs that were not annotated correctly. Two sequences from *Lymnaea stagnalis* were also included as they were functional characterized as anionic and cationic ACh-gated channels^23^. *Platynereis dumerilli (GCA_026936325.1)*, *Hellobdella robusta (Helro1)*, and *Capitella teleta (GCA_000328365.1)* were included as annelid representatives. Three sequences from *Alvinella pompejana* were also included as they had been previously functionally identified as glycine and proton-gated channels^35^. *Schmidtea mediterranea* (PRJNA379262) and *Schistosoma mansoni* (PRJEA36577) were included for Platyhelminths. *Drosophila melanogaster (BDGP6.46)*, *Daphnia magna (ASM2063170v1.1)*, and *Apis mellifera* (Amel_HAv3.1) were included for Arthropods. *Caenorhabditis elegans* (PRJNA13758) and *Brugia malayi* (PRJNA10729) were included for nematodes.

*Nematostella vectensis (PRJEB51373)*, and *Clytia hemisphaerica* ^36^ represented the Cnidarians. All sequences were downloaded from Biomart ^37^, except for *O. vulgaris* where we ran interproscan (v5.33) ourselves and *C. hemisphaerica* where we downloaded the sequences from MARIMBA (http://marimba.obs-vlfr.fr/home). Jaiteh et al. ^38^ determined that Ctenophora, Proifera and Placozoa had no LGCs in their genomes. We also included all metazoan sequences from the Jaiteh et al., 2016 dataset, which included 31 representative metazoan species.

#### Alignment and tree generation

LGCs have four transmembrane domains, and thus partial sequences with less than three transmembrane domains were removed. Transmembrane domains were predicted using TMHMM ^39^. The longest isoform for each gene was manually identified and others were removed. Alignment was performed using MAFFT (v7.505) with E-INS-i parameters allowing for large gaps ^40^, as Jaiteh et al., ^38^, determined this was an effective approach for aligning LGC protein sequences. The alignment was trimmed using Trimal (v1.4.1) gappyout method ^41^. The alignment was manually assessed to ensure each sequence had the ‘cys-loop region’, and the four transmembrane domains. Any sequences with large inserts were removed iteratively until a satisfactory alignment was achieved.

Phylogenies were generated using IQ-TREE v2.1.4_beta with 1,000 ultra-fast bootstraps ^42^. The tree model was calculated using IQ-TREE’s ModelFinder implementation according to Bayesian inference criterion (BIC) and Q.yeast+F+R10 was used.

#### Tree visualization and annotation

The tree was visualized using TreeViewer ^43^ and species illustrations were either generated by authors or sourced from Phylopic (phylopic.org) (Fig. 2 a and e). LGCs contain a characteristic ‘cys-loop’ region in their extracellular ligand binding domain which contains 13 residues flanked by two cysteines that create a disulphide bond. Jaiteh et al. previously characterised a ‘cys-less’ group of LGCs which lack these cysteines in eukaryotes (many protists and a few invertebrates) ^38^. We used this cys-less group to root the tree. We assigned a functional annotation for the primary ligand to any subfamily where one or more sequences had been previously functionally characterized in a heterologous expression system. Our LGC tree, and others in the literature, split between GABA-A-like anionic LGCs and Nicotinic-ACh-like cationic LGCs. However, within these anionic/cationic subfamilies some exception exists and thus the ion selectivity of these exceptional cases are also highlighted where functional data exists. The references associated with the functional annotations of each sequence can be found in supplemental file 2. Our LGC dataset contains 1197 sequences, the alignment is available as supplemental file 3 and the full tree is available as supplemental file 4.

### Two-electrode voltage clamp recordings of LGCs in *Xenopus* oocytes

#### *Xenopus* laevis plasmids

*O. vulgaris* RNA was extracted from 15 1-dph paralarvae using Tri-reagent (Invitrogen) and the Qiagen Micro kit (Qiagen). *Daphnia magna* RNA was extracted from 20 adults using Tri-reagent (Invitrogen) and the Qiagen Micro kit (Qiagen). cDNA for both *O. vulgaris* and *D. magna* was synthesised using the Superscript III RT (Invitrogen) kit, and LGC sequences were amplified using Q5 polymerase (NEB, MA, USA). The cDNA sequence for the DopC1 gene from *Aplysia californica* (LOC101853763) and *Apis mellifera* (LOC408525) were synthesised using the GeneArt Gene Synthesis Service (Thermo Fisher Scientific, Waltham, MA, USA). LGC cDNA sequences from *O. vulgaris, D. magna, A. californica and A. mellifera* were then inserted into the KSM backbone using HiFi assembly (NEB, MA, USA). The KSM backbone contains *Xenopus* β-globin UTR regions and a T3 promoter. The cDNA sequence of the *Octopus bimaculoides* AChRB1 (LOC106879694/XM_052969792.1) gene was synthesized as a geneblock DNA fragment by IDT-DNA Inc (Coralville, IA, USA) with a Kozak transcriptional start signal (GCCACC) added immediately upstream of the initial methionine codon. This fragment was inserted between the two Sma I restriction enzyme sites of pGEMHE ^44^ by Gibson assembly.

#### *Xenopus laevis* oocytes and RNA injection

Defolliculated *Xenopus Laevis* oocytes were acquired from EcoCyte Bioscience and maintained in ND96 solution (in mM: 96 NaCl, 1 MgCl2, 5 HEPES, 1.8 CaCl2, 2 KCl) at 16°C until RNA injection. KSM plasmids were linearized with NotI, and 5’-capped cRNA was synthesized *in vitro* using the T3 mMessage mMachine transcription kit (Thermo Fisher Scientific, CA, USA). The pGEMHE plasmid was linearised with Eco53kI and synthesized *in vitro* using the T7 mMessage mMachine transcription kit (Thermo Fisher Scientific, CA, USA). The RNA was subsequently purified with the GeneJET RNA purification kit (Thermo Fisher Scientific, CA, USA). Defolliculated *Xenopus* oocytes were individually placed in v-bottom 96-well plates and injected with RNA using the Roboinject system (Multi Channel Systems GmbH, Reutlingen, Germany). Each oocyte was injected with 50 nl RNA solution at a concentration of 250 ng/µl-500 ng/µl, depending on the construct. Following injection, the oocytes were incubated in ND96 at 16°C for 2-4 days prior to recording.

#### Two-electrode voltage clamp recording and analysis

TEVC recordings were performed either using the automated Robocyte2 system (Multi Channel Systems, Reutlingen, Germany) or a manual system (OC-725D amplifier). Recording electrodes for the Robocyte2 system were purchased from Multichannel systems and typically had a resistance of 0.7-2 MΩ. Recording electrodes for the manual system (GC120TF-10, Warner Instruments) were pulled using a P-1000 Micropipette Puller (Sutter, Ca, USA) with the same resistance as above. Recording electrodes were filled with 1.5M KCl and 1 M acetic acid. Oocytes were clamped at -60 mV and continuous recordings were done at 500 Hz. Recordings on the Robocyte2 system were collected using the Robocyte2 control software and the data was extracted using the Robocyte2+ analysis software.

Recording on the manual system were collected using WinWCP software. The following ligands were applied to all receptors described in the paper at 1mM during initial experiments to determine their primary/secondary ligands. All ligands were prepared in ND96 solution and were applied for 7 seconds. Fresh ND96 solution then applied for 40s to wash out the ligand before applying the next ligand. The ligands used for screening were acetylcholine chloride (ACh, sc-202904, Santa Cruz Biotechnology,USA), L-Glutamic acid monosodium salt monohydrate (Glut, 49621, Merck, Germany), γ-Aminobutyric acid (GABA, A2129, Merck, Germany), Glycine hydrochloride (Gly, G2879, Merck, Germany), dopamine hydrochloride (Dop, H8502, Merck, Germany), DL-norepinephrine hydrochloride (Norepi, A7256, Merck, Germany), (+/-)-Epinephrine hydrochloride (Epi, E4642, Merck, Germany), Tyramine hydrochloride (Tyra, T2879, , Merck, Germany), (+/-)-Octopamine hydrochloride (Octo, O0250, Merck, Germany), serotonin hydrochloride (5-HT, sc-201146A, Santa Cruz Biotechnology, USA), Choline chloride (Chol, C1879, Merck, Germany), Betaine hydrochloride (Bet, B3501, Merck, Germany), adenosine (Ado, 01890, Merck, Germany), Adenosine 5′-triphosphate disodium salt hydrate (ATP, A26209, Merck, Germany), β-alanine (β-al, 146064, Merck, Germany), D-serine (D-ser, S4250, Merck, Germany), and L-Lysine (Lys, L5501, Merck, Germany).

Dose response analysis was performed by first perfusing the oocytes with ND96 to estimate resting potential of the oocytes. Then the ligand solution was applied for 7 seconds and then ND96 solution was applied again for 40 seconds to bring the oocyte back to resting potential. This was repeated for all doses of the ligand from low to high concentrations. The dose response results combined data from 3-7 oocytes and the data was normalised to the largest current in each oocyte. Normalized mean and standard deviation where then imported into GraphPad prism where data was plotted and EC_50_ values were calculated by fitting to the Hill equation using three parameter slopes to obtain the highest degree of fit.

Ion selectivity analysis was performed using ND96 solution, a sodium-free solution (96mM NMDG, 2mM KCl, 1.8mM CaCl2, 5mM HEPES, 1mM MgCl2) and a low-chloride solution (96mM Na Gluconate, 2mM KCl, 1.8mM CaCl2, 5mM HEPES, 1mM MgCl2). A voltage ramp from -150 mV to +30 mV was applied to oocytes expressing the receptor of interest in the presence of the following solutions in the following order; ND96, ND96+ligand, sodium-free, sodium-free +ligand, low-chloride, low-chloride +ligand. The EC_50_ concentration of the primary ligand was used in each case, either ACh for AChRBs and dopamine for DopCs. Analysis was performed using a custom python script (https://github.com/hiris25/TEVC-analysis-scripts).

Peak current analysis was performed by exposing individual oocytes expressing specific LGCs with 1mM dopamine, serotonin, norepinephrine, octopamine, epinephrine, and tyramine in this order and the current was normalized to the peak current in each oocyte. After exposing each oocyte to these ligands, dopamine was applied again at 1mM to ensure the channel had not desensitized. The mean and standard deviation per channel and per ligand were then calculated and the data was plotted in Graphpad Prism.

### Single cell RNA sequencing analysis

All scRNA-seq results were from the dataset detailed in Styfhals et al ^9^. Data visualisation was performed in Seurat v4 ^45^.

### RNA fluorescent *in situ* hybridisation chain reaction (HCR)

1-dph paralarvae were sedated in 2% ETOH/artificial seawater, humanely killed in 4% EtOH/artificial seawater and fixed overnight in 4% paraformaldehyde (PFA) in phosphate-buffered saline treated with DEPC (PBS-DEPC). RNA fluorescent *in situ* hybridization chain reaction (HCR) was performed as previously described ^46^. In brief, parlarvae were embedded in paraffin following progressive dehydration and sectioned with a paraffin microtome (Thermo Scientific, Microm HM360) to obtain 6-µm-thick transversal sections. Prepared tissue slides were then used to perform hybridization chain reaction experiments (HCRv3.0). Probe sets were designed with the insitu_probe_generator ^47^, followed by automated blasting and formatting to minimize off-target hybridization with a custom script ^48^ and synthesised by IDT (DNA oPools™, Integrated DNA Technologies). Probes were blasted against the *O. sinensis* (PRJNA551489) or the *O. vulgaris* transcriptome (PRJEB61268) ^34^ to ensure specificity. Each probe was tested on multiple occasions on different batches of paralarvae and the expression pattern reproducible. These probes were applied at 0.9 pmol per slide. Upon completion of the HCR protocol, DAPI staining was performed (32670-5MG-F, Merck), slides were mounted with a cover slip (Epredia, 24 x 55 m #1) using Mowiol and allowed to dry for 24 h at 18°C before being stored at 4°C until imaging. All probes sequences are included in supplemental file 5.

#### HCR image acquisition

Confocal microscopy was performed using a spinning disk system (Andor BC43, Oxford Instruments) with 405 nm, 562 nm, 600 nm, and 700 nm excitation lasers (respectively blue, green, red and far red). Images were acquired with a 60X/1.42 NA oil-immersion objective (Nikon CFI Plan Apochromat Lambda D) and an sCMOS camera (Oxford Instruments, 6.5 µm pixel size, 82% QE) at a 0.3 µm z-step.

#### HCR image analysis

All HCR images were prepared for publication by cropping and adjusting brightness/contrast using Imaris Software (Oxford Instruments) and Fiji ^49^. HCR fluorescence was not visible in the plexiform layer or other neuropil regions and localised around the nuclei, therefore a custom python script was written to quantify HCR fluorescence within the nuclear regions of cells in the *O. vulgaris* paralarval brain (https://github.com/jboulanger/fish-octopus). The expression pattern was consistent across replicates and thus quantification was performed on individual slices. The cell pose pre-trained ‘Nuclei’ model was used to segment nuclei (DAPI channel) in 3D from the Z-stack projections^50^. Each 6µm section contained ∼2 layers of densely packed nuclei. The Napari polygon tool was used to manually define anatomical regions (i.e. OGL, IGL, Med, CB) (https://github.com/napari/napari). A difference of Gaussian filter with sigma [0.5,1,1 pixels] along the z,y,x axes was used to identify positive fluorescent signals from background signal. The filtering was followed by a Relu activation defined as max 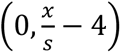 where the standard deviation *s* is estimated using a maximum absolute deviation to define a score. The mean fluorescence of this score for each fluorescent channel within the area of the 3D segmented nuclei was recorded. A second threshold was then manually defined for each gene to determine positive/negative expression within a segmented nucleus.

### Calcium imaging of acute paralarval brain slices

#### Reagents and solutions

Octomedia was prepared by starting with 500 ml L-15, Leibowitz, with L-glutamine (Sigma, MFCD00217482) and then adding the following components (in mM) NaCl: 214.04, MgSO_4_x7H_2_O: 26.17, KCl: 4.61, NaHCO3: 2.29, MgCl_2_x6H_2_O: 28.04 , D-glucose: 38.08, and CaCl_2_x2H_2_O: 10.12. Taking the composition of L-15 into account the final concentrations were; (in mM) NaCl: 351.97, MgSO_4_x7H_2_O: 26.984, KCl: 9.94, NaHCO3: 2.29, MgCl_2_x6H_2_O: 29.026, D-glucose: 38.08, CaCl_2_x2H_2_O: 11.38, C3H3NaO3: 5, KH₂PO₄: 0.44, Na_2_HP0_4_: 1.34. HighK^+^ Octomedia is identical to Octomedia except NaCl and KCl were added as 168.65 mM and 50 mM respectively, which yielded a final concentration of 306.58 mM and 55.33 mM. Both solutions were adjusted to pH 7.6-7.7 with KOH, filtered using the corning system (431098), stored at 4°C and used within one week. Calcium indicator CAL-520 (Abcam, UK) was prepared at 5 mM in DMSO, and stored at -20°C. Dopamine, and ACh were prepared at 100 µM in Octomedia. Agarose (11AGAL0050, MP Biomedicals) with high gel strength (500 g/cm²) and low gelling temperature (26°C) was prepared at 3% in Octomedia and kept in a 42°C water bath until ready to use.

#### Brain dissection, embedding and sectioning

1-dph paralarvae were anaesthetised in 2% ETOH/filtered artificial seawater (FASW:) until motionless (at least 5 minutes). Their brains were dissected from their bodies to ensure all surrounding skin/muscle were removed. The brain was then placed in the 3% agarose solution in a 35mm petri dish. The agarose was left to solidify at room temperature for 5 minutes and then stored on ice until vibratome sectioning (within an hour). The agarose was cut to a 1cm x 1cm block and sectioned with a vibrating-blade microtome (HM 650 V, Thermo Scientific, UK) with the following parameters; 200 µm slice thickness, 0.2 mm/s speed, and amplitude 0.7mm. The vibratome bath was filled with cooled Octomedia. Brain slices were transferred to 25mm diameter circular coverslips. A temporary well was created using a plastic 3D printed cylinder made from the plastic polylactic acid (10mm height, 3mm wall thickness, 18mm diameter, supplemental file 6), and petroleum jelly was placed on the rim of the cylinder to create a seal to the coverslip. Each slice was then incubated in CAL-520 (5µM in Octomedia) for at least 1 hour at room temperature within this temporary well. The cylinder was then removed and the circular coverslip containing the slice was directly transferred to the imaging chamber.

#### Calcium imaging and perfusion

Imaging was performed with an upright epifluorescent microscope (Leica DM6 B) equipped with a 490/525 nm excitation/emission filter set and operated at 4-5fps, which was limited by exposure time (200-250 ms). The light intensity was as low as possible, dictated by a manual knob on the light source. The iris was also set as low as possible to further reduce scattered light and photobleaching. Perfusion inflow and compound switching were controlled by a gravity valve system (Digitimer, UK), while outflow was maintained with a peristaltic pump. The imaging chamber (PH2 and RC-21BDW, Science Products) was assembled with 22×40 mm rectangular coverslips creating a lid. The slices were immobilised using netting (hexagonal pore size: 1mm x 1.5 mm) that lay on the agarose surrounding the brain slice.

Slices were perfused with Octomedia for 10 minutes before imaging to acclimatise the slice to the imaging chamber. The perfusion protocol during imaging was as follows; Octomedia: 0-90s, Ligand: 90s-135s, Octomedia: 135s-270s, HighK^+^ Octomedia: 270s-end:

### Calcium imaging analysis

A total of 973 cells across 17 slices were analysed during dopaminergic stimulation in the OGL of the OL. Likewise, 1090 cells over 18 slices and 2298 cells over 18 slices were analysed in the IGL and medulla respectively. In terms of ACh stimulation, we analysed 391 cells across 12 slices from the OGL, 473 cells across 12 slices from the IGL and 858 cells across 12 slices from the medulla. The calcium imaging recordings of each slice were processed as follows using the Suite2p GUI^51^ and a custom python scripthttps://github.com/A-Courtney/Octo_Dop_ACh_Ca_Imaging_2025).

#### Image registration

We observed deformations of the tissue slice during recording sessions, likely due to muscular contractions, particularly during application of the HighK^+^ solution. We assume that they are caused by the oesophagus that runs through the centre of the brain and perhaps muscle fibres surrounding the brain that were insufficiently removed during dissection. To overcome this, image registration was performed within suite2p^51^ using the non-rigid registration approach and 128 x 128 pixels block size (https://github.com/MouseLand/suite2p).

#### ROI detection

Regions of interest (ROIs), corresponding to cells, were detected within suite2p using an anatomical detection approach. This was achieved in a semi-automated way by applying the cell pose ‘cyto3’ model to the maximum intensity projection image of each slice^50^, and the ROIs were then refined manually. Then within the suite2p GUI any cells that were incorrect were deleted and any cells that were missing were added manually. Out of focus regions were not analysed.

#### ROI quality control

The ROIs detected using the anatomical approach described above were a combination of ‘real’ cells, ‘dead’ cells and ‘neuropil’ (fluorescence signals from local neurites). Therefore, we wanted to remove the ‘dead’ and ‘neuropil’ ROIs and only analyse the ‘real’ cell ROIs. We first wanted to understand the functional characteristics of ‘cellular’ ROIs vs ‘neuropil’ ROIs, we manually defined multiple ROIs using fiji^49^ from five slices from different paralarvae exposed to both dopamine and ACh (Supplemental Fig. 15 a-d). Manually defined ‘cellular’ ROIs had a higher fluorescence and a circular structure. Manually defined ‘neuropil’ ROIs had a lower fluorescent signal and were found in between the ‘cellular’ ROIs. We calculated the slopes in two second windows across the full normalised fluorescent trace. Slope was calculated by using the difference in fluorescence between the start and end of the time window, divided by the number of frames. We observed that ‘cellular’ ROIs had a statistically significantly higher max slope compared to ‘neuropil’ ROIs (t-test, p<0.05, Supplemental Fig. 15 e). We then used the mean+standard deviation of the neuropil max slope (0.0218 normalised intensity/second) as a threshold so all traces that had a max slope below this are likely a heavily neuropil contaminated ROI and were removed from subsequent analysis. This threshold also filtered out ‘dead’ ROIs as the traces from these ROIs have a downward fluorescent signal over the course of the recording due to photobleaching and don’t show the same characteristic responses to the ligand, HighK^+^ solution or any spontaneous activity.

#### Defining anatomical regions

To analyse the functional characteristics of cells within different regions of the optic lobe of *O. vulgaris* paralarvae, we used Napari (https://github.com/napari/napari) to manually define anatomical regions such as the OGL, IGL, medulla of the OL. The maximum intensity projection of each slice was used to label these regions with the polygon tool.

#### Ligand response categorisation

The ligand response for each trace was defined as either excitation, inhibition or no-response. Excitation was defined as the ligand stimulus window having a 20% higher mean fluorescence compared to the baseline mean fluorescence (first 20 seconds of the recording), when the full trace, including the HighK^+^ response, was normalised using a min/max approach (Supplemental Fig. 15 f shows example excitatory responses following dopamine application. Inhibitory signals had to meet multiple criteria; 1. the ligand stimulus window had to have a 20% decrease in mean fluorescence compared to the baseline mean value (first 20 seconds of the recording), when the full trace, including the HighK^+^ response, was normalised using a min/max approach, 2. inhibitory responses also needed to have a steep downward slope (<-0.0099 normalised intensity/s) and, 3. the trough of the inhibition needed to last at least 20 seconds. Supplemental Fig15 g shows example inhibitory responses following ACh application. ROIs with ‘no response’ didn’t meet either the excitation or inhibition criteria and usually had a flat response during ligand application and a normal HighK^+^ response (Supplemental Fig. 15 f and g). Subtle differences in perfusion speed are inevitable on different experimental days, and thus ligand and HighK^+^ response windows were manually defined.

#### Dopamine slope analysis

All dopaminergic excitatory responses that were identified above were then subsequently analysed for their slope. Within each ROI trace the dopaminergic response had different onset times, so the time window for calculating the dopamine response slope could not be fixed.

Instead, the dopamine application time window trace was extracted (80 second window), smoothed using a rolling average over adjacent frames (4 second window), and normalized using the min/max values. The slope is calculated as above, the time window was 2 seconds and the slope for all 2 second periods across the stimulus time window were calculated, and the max slope reliably identified the steepest slope of the dopaminergic response for each ROI.

Statistical analysis of the slope of excitatory dopaminergic responses involved first checking for normality per region (OGL, IGL and Med) using a Shapiro-Wilk test and none of the groups were normally distributed (p < 0.05). The variances between the regions were not equal (Levene’s Test, p <0.05) and thus Welch’s ANOVA was used to determine if there was a significant difference between the regions. Games-Howell post-hoc test was used to determine which optic lobe layers had statistically significant differences in their dopamine slope.

While only ‘in-focus’ cells were detected and analysed, there were still some variations in how ‘in-focus’ a cell was. To ensure that this didn’t impact our slope calculations we first looked at the radius of cells, assuming cells with a larger radius were more out of focus, and compared it to the slope of excitatory dopaminergic response. We performed Spearman correlation and we did not observe a correlation between ROI radius and slope (R^2^=-0.037, p=0.0482, Supplemental Fig. 15 h). We also used variance of laplacian to calculate the ‘sharpness’ of a 30 by 30-pixel window surrounding the ROI ^52^ and compared it to the slope of excitatory dopaminergic response. We did not observe a correlation between ROI sharpness and slope (Spearman correlation, R^2^=0.009, p=0.5647, Supplemental Fig. 15 i).

---

**Supp Fig. 1.**
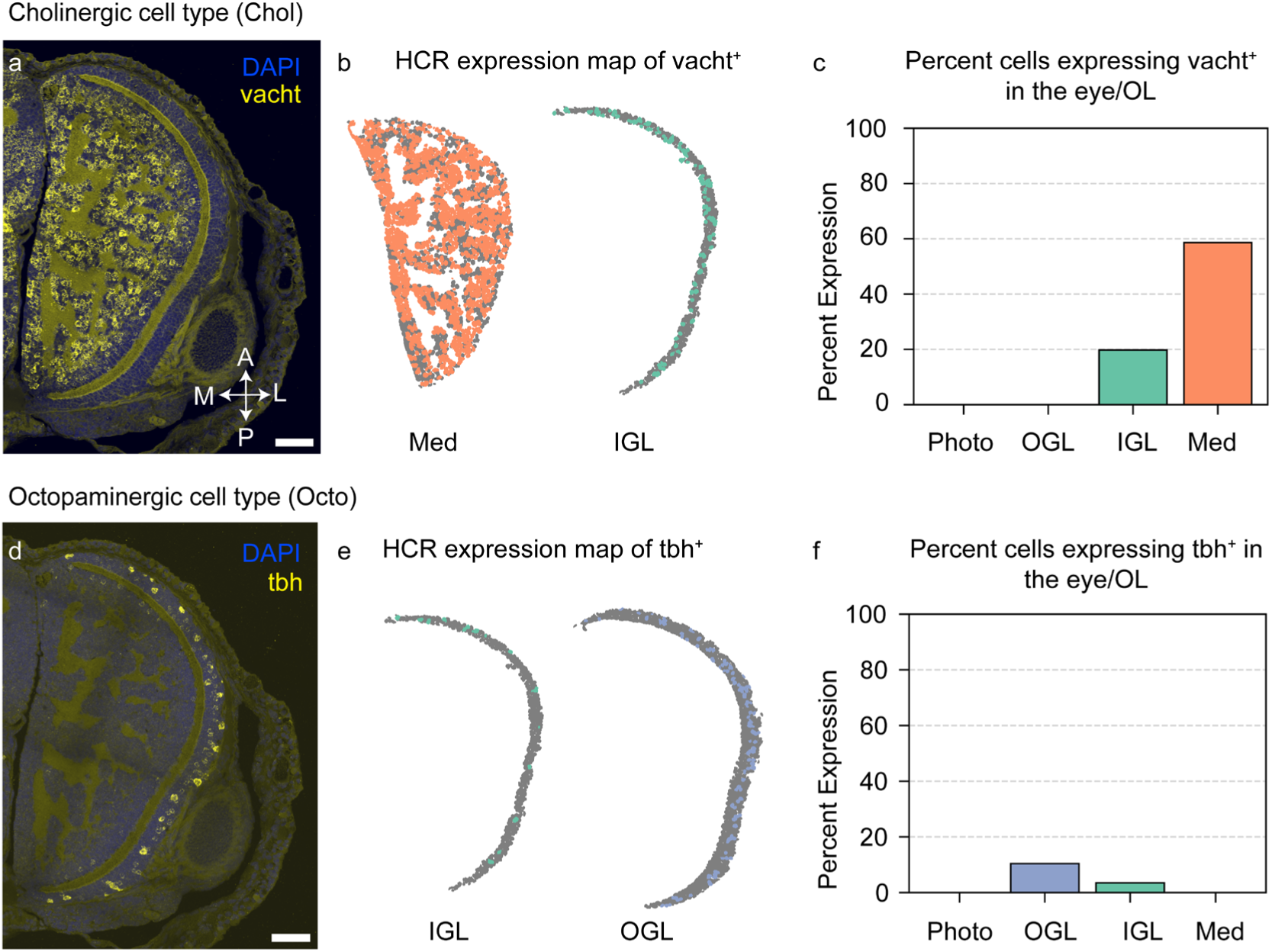
Quantitative and spatial analysis of Chol and Octo cell types in the visual system of 1-dph O. vulgaris paralarvae. HCRs were performed on 1-dph *O. vulgaris* head transversal sections and nuclei are visualised in blue with DAPI. (**A**). Representative HCR against vacht (cholinergic cell marker) in yellow. A custom python script was used to quantify the expression pattern for each gene of interest within nuclear regions in the eye and optic lobe (see methods). (**B-D**) HCR expression maps (**B**) and barplot (**C**) showing the proportion and spatial distribution of cells expressing vacht in the eye/OL, quantified from the HCR shown in A. (**D**). Representative HCR against tbh (octopaminergic cell marker) in yellow (**E-F**) HCR expression maps (**E**) and barplot (**F**) showing the proportion and spatial distribution of cells expressing tbh in the eye/OL, quantified from the HCR shown in D. The orientation in A and D are the same. Scale bar: 50 μm, vacht: vesicular acetylcholine transporter, tbh: tyramine beta hydroxylase, Photo: photoreceptors, OGL: outer granular layer, IGL: inner granular layer, Med: medulla, OL: optic lobe, Octo: octopaminergic cell type (tbh^+^), Chol: cholinergic cell type (vacht^+^), A: anterior, P: posterior, M: medial, L: lateral.

**Supp Fig. 2.**
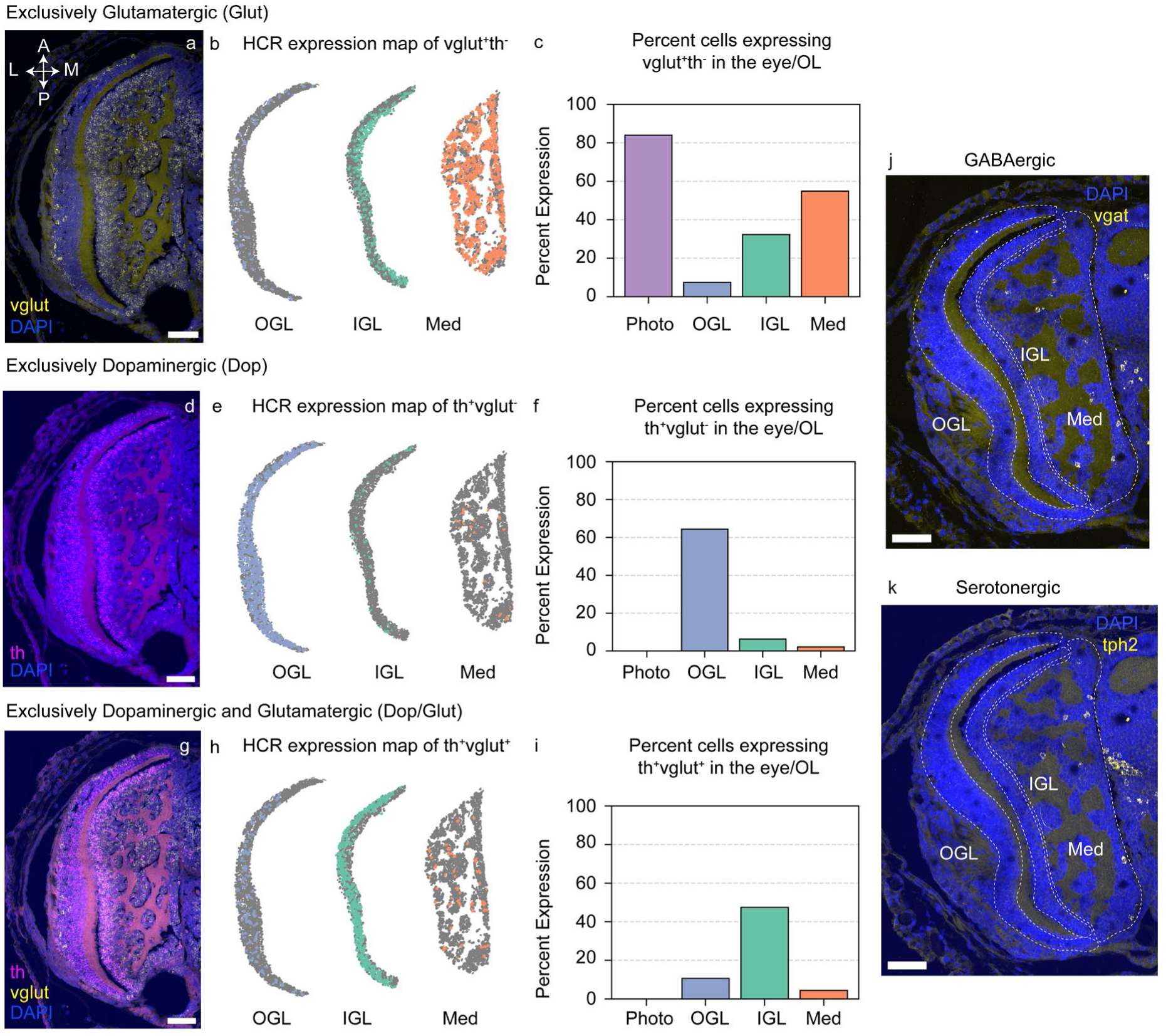
Quantitative and spatial analysis of Glut, Dop and Dop/Glut cell types in the visual system. All HCRs were performed on 1-dph *O. vulgaris* head transversal sections; nuclei were visualised in blue with DAPI. (**A**) Representative HCR against the glutamatergic marker vglut (vesicular glutamate transporter) is shown in yellow. A custom python script was used to quantify the expression pattern for each gene of interest within nuclear regions in the eye and optic lobe (see methods). HCR expression maps (**B**) and barplot (**C**) show the proportion and spatial distribution of cells expressing vglut exclusively (vglut^+^th^−^) in the eye/OL, quantified from the HCR shown in A. (**D**) Representative HCR against dopaminergic marker th (tyrosine hydroxylase) in magenta. HCR expression maps (**E**) and barplot (**F**) show the proportion and spatial distribution of cells expressing th exclusively (th^+^vglut^−^) in the eye/OL, quantified from the HCR shown in D. (**G**) Representative HCR against both th (in magenta) and vglut (in yellow). HCR expression maps (**H**) and barplot (**I**) show the proportion of Dop/Glut cells (th^+^vglut^+^) in the eye/OL, quantified from the HCR shown in G. (**I-K**) Representative HCR against GABAergic marker vgat (vesicular GABA transporter) (**J**) and serotonergic cell marker tph2 (tryptophan hydroxylase) (**K**) in yellow. All HCRs have the same orientation. Scale bar: 50 μm, vgat:, tph2:, th: tyrosine hydroxylase, vglut:, Photo: photoreceptors, OGL: outer granular layer, IGL: inner granular layer, Med: medulla, OL: optic lobe, Dop: exclusively dopaminergic cell type (th^+^vglut^−^), Glut: exclusively glutamatergic cell type (vglut^+^/th^−^), Dop/Glut: dopaminergic and glutamatergic cell type (th^+^vglut^+^), A: anterior, P: posterior, M: medial, L: lateral.

**Supp Fig. 3.**
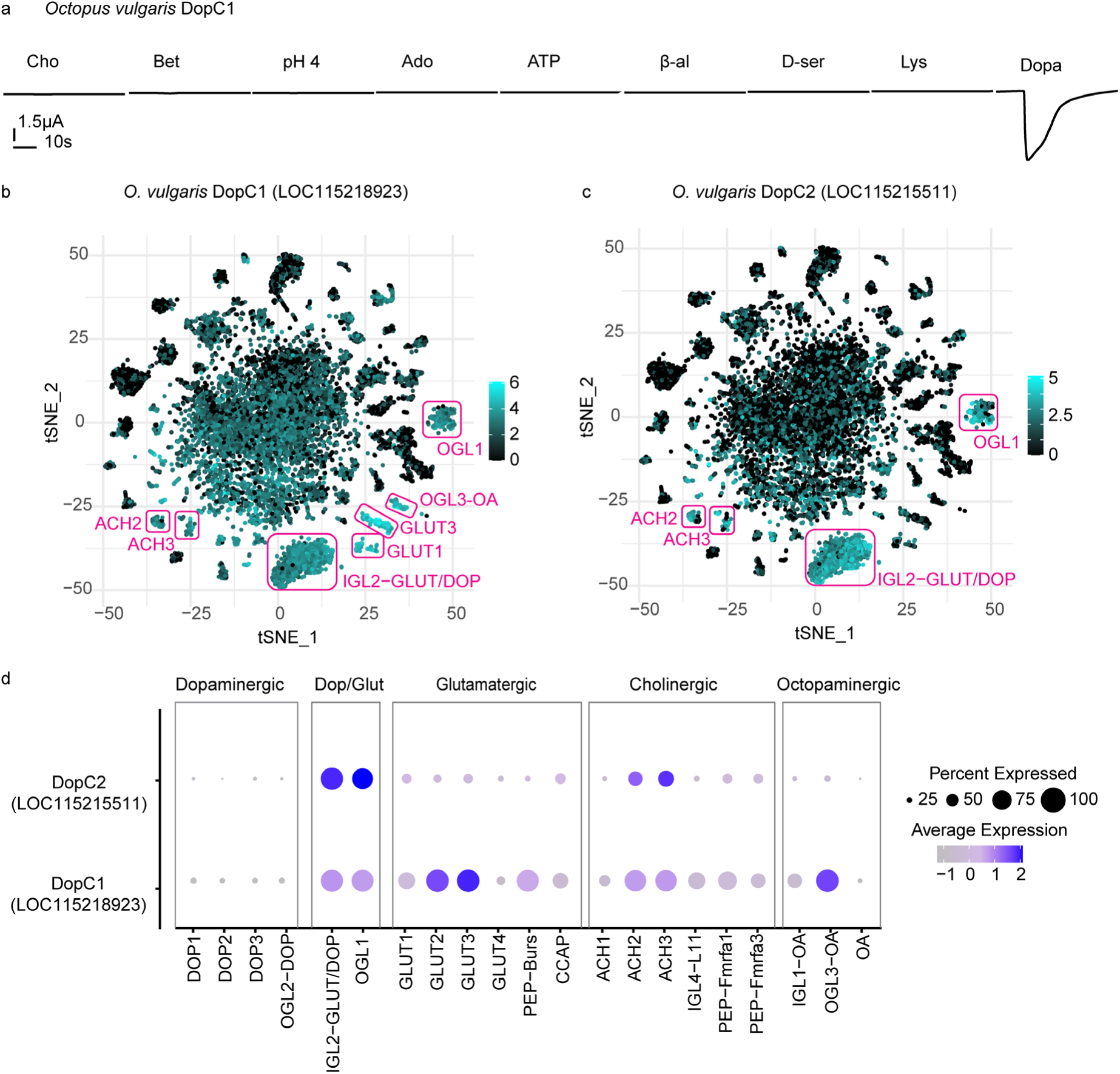
Additional functional and expression analysis of DopC1 from *O. vulgaris*. (**A**) Representative TEVC traces from *Xenopus* oocytes expressing DopC1 from *O. vulgaris* during application of additional candidate ligands. (**B-C**) tSNE of scRNA-seq of the *O. vulgaris* paralarval brain^9^, showing the expression of the DopC1 (LOC115218923) (**B**) and DopC2 (LOC115215511) (**C**). All clusters that show high expression are highlighted in magenta. (**D**) Dot plot of a subset of clusters from the single cell RNAseq of the *O. vulgaris* paralarval brain showing the expression of DopC1 and DopC2. The size of the dots relates to the percent of cells in that cluster that express the gene, the heat map corresponds to the average expression of the gene in all cells in the cluster. Cho: choline, Bet: betaine, Ado: adenosine, ATP: adenosine triphosphate, β-al: β-alanine, Lys: lysine, Dopa: dopamine.

**Supp Fig. 4.**
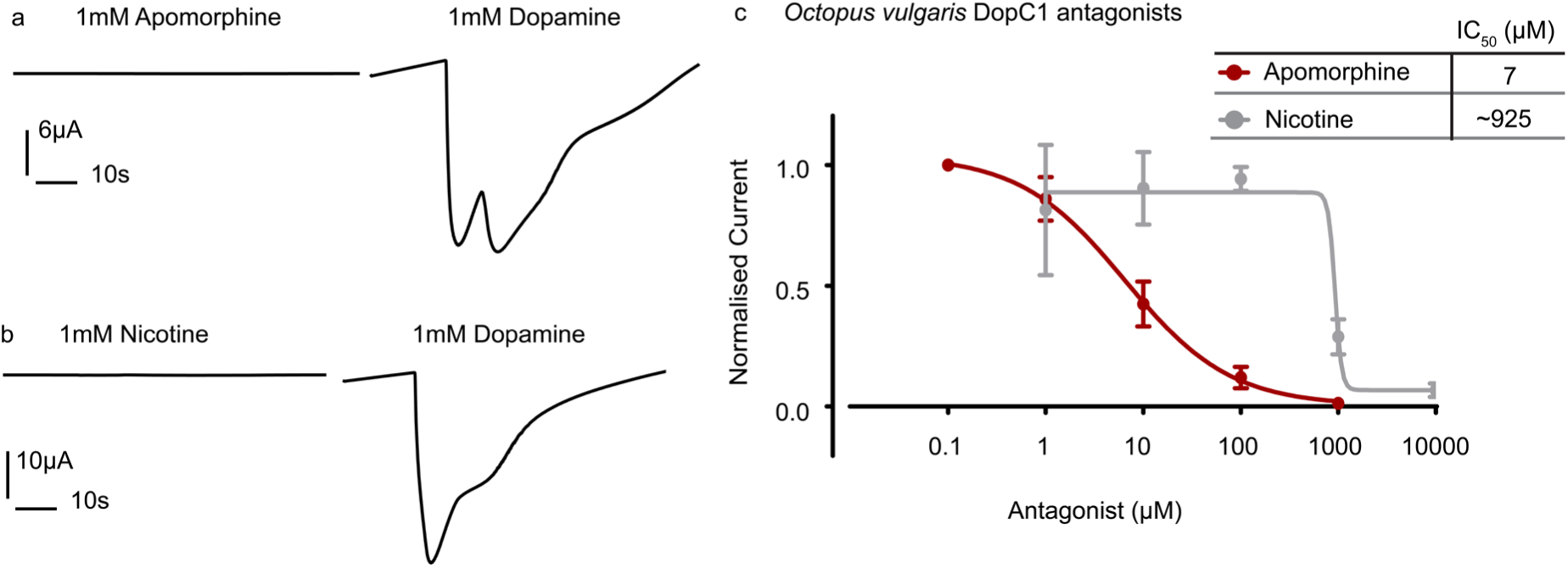
Characterising Octopus vulgaris DopC1 candidate agonists and antagonists. (**A-B**) Representative TEVC traces from *Xenopus* oocytes expressing *O. vulgaris* DopC1 during application of apomorphine (**A**) and nicotine (**B**). (**C**) Antagonist inhibitory dose response curves from *Xenopus* oocytes expressing DopC1 from *O. vulgaris*. This was acquired by applying low to high doses of each antagonist in the presence of dopamine at 10µM (∼EC_50_). The adjacent table includes the IC_50_ values for each ligand in µM. Mean and standard error are plotted for 5-7 independent oocytes per antagonist. The current was normalised to the max current in each oocyte.

**Supp Fig. 5.**
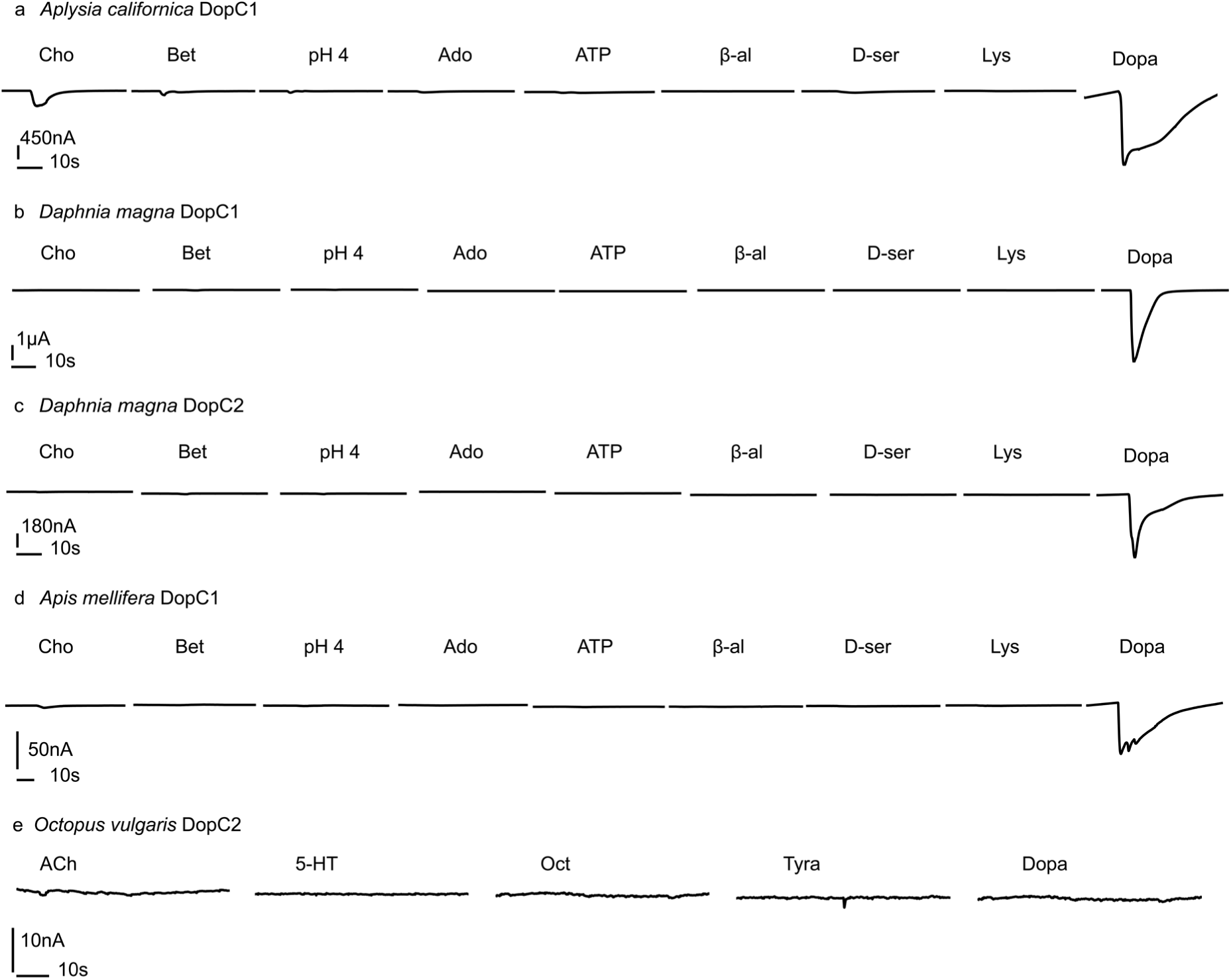
Functinal characterization of DopC orthologs from *Aplysia californica, Apis mellifera* and *Daphnia magna*. Representative TEVC traces from *Xenopus* oocytes expressing DopC1 from *Aplysia californica* (**A**), DopC1 from *Daphnia magna* (**B**), DopC2 from *Daphnia magna* (**C**), DopC1 from *Apis mellifera* (**D**) and DopC2 from *Octopus vulgaris* (**E**) during application of candidate ligands. ACh: acetylcholine, Cho: choline, Bet: betaine, Ado: adenosine, ATP: adenosine triphosphate, β-al: β-alanine, D-ser: D-serine, Lys: lysine, Dopa: dopamine, Oct: octopamine, Tyra: tyramine, 5-HT: serotonin, DopC: dopamine-gated cation channel.

**Supp Fig. 6.**
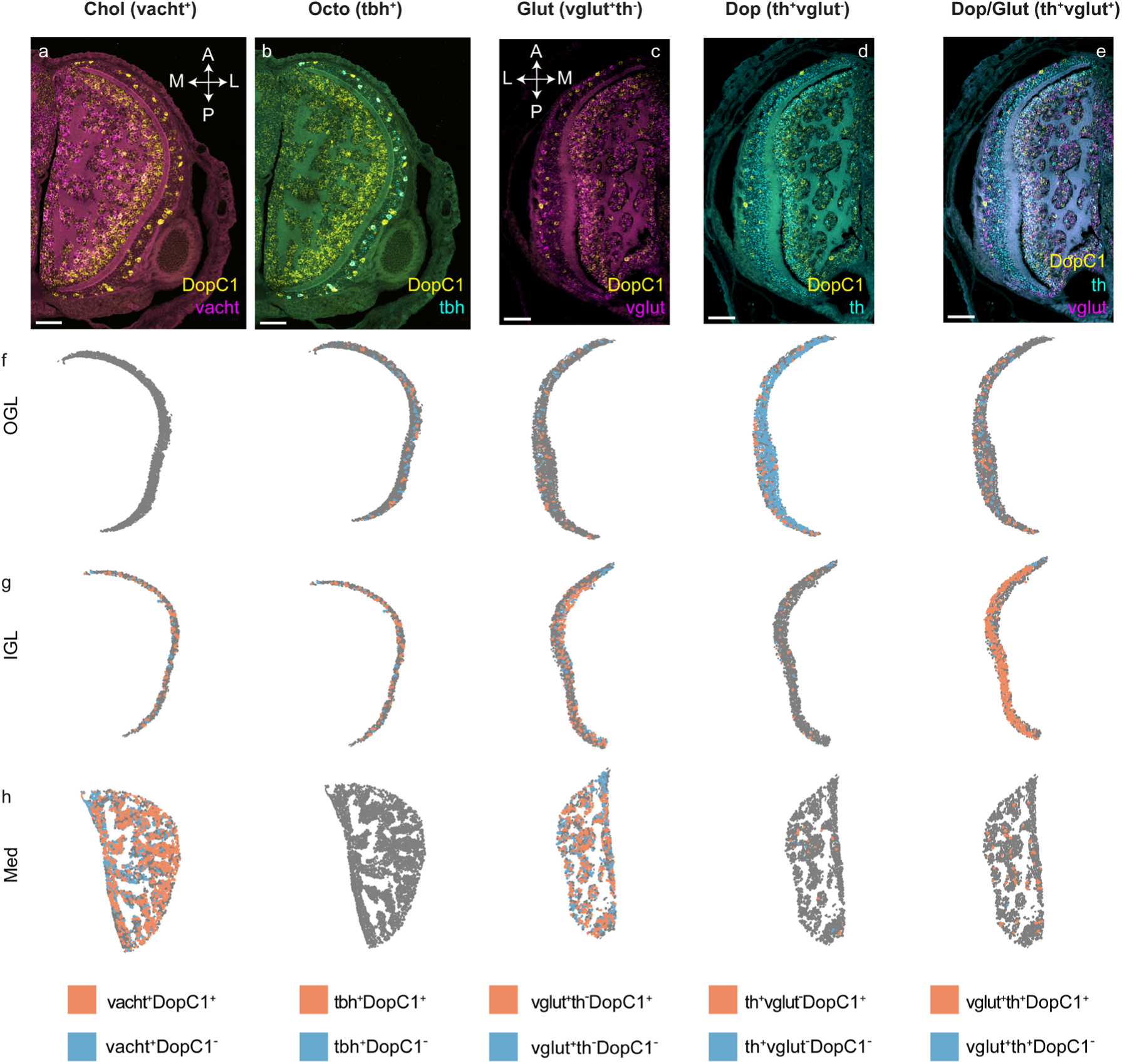
Quantitative and spatial analysis of DopC1 expression in the brain of *O. vulgaris*. Representative multiplex HCR against DopC1 in yellow and vacht (cholinergic marker) in magenta (**A**), tbh (octopaminergic marker) in cyan (**B**), vglut (glutamatergic marker) in magenta (**C**) and th (dopaminergic marker) in cyan (**D**). (**C** and **E**) Representative multiplex HCR against DopC1 in yellow, vglut (glutamatergic cell marker) in magenta (**C**) and th (dopaminergic cell marker) in cyan (**E**). (**F-H**) Representative HCR expression maps of Chol, Octo, Glut, Dop and Dop/Glut cell types in the OGL (**F**), IGL (**G**) and Med (**H**) of the OL, with cells positive for DopC1 in orange and those negative for DopC1 in blue. The expression maps were quantified from the HCRs shown above and the results are summarised in Fig. 4 i. The orientation in A and B and C-D are the same. Cell type, layer, and gene names are as described in previous figures. Scale bar: 50 µm.

**Supp Fig. 7.**
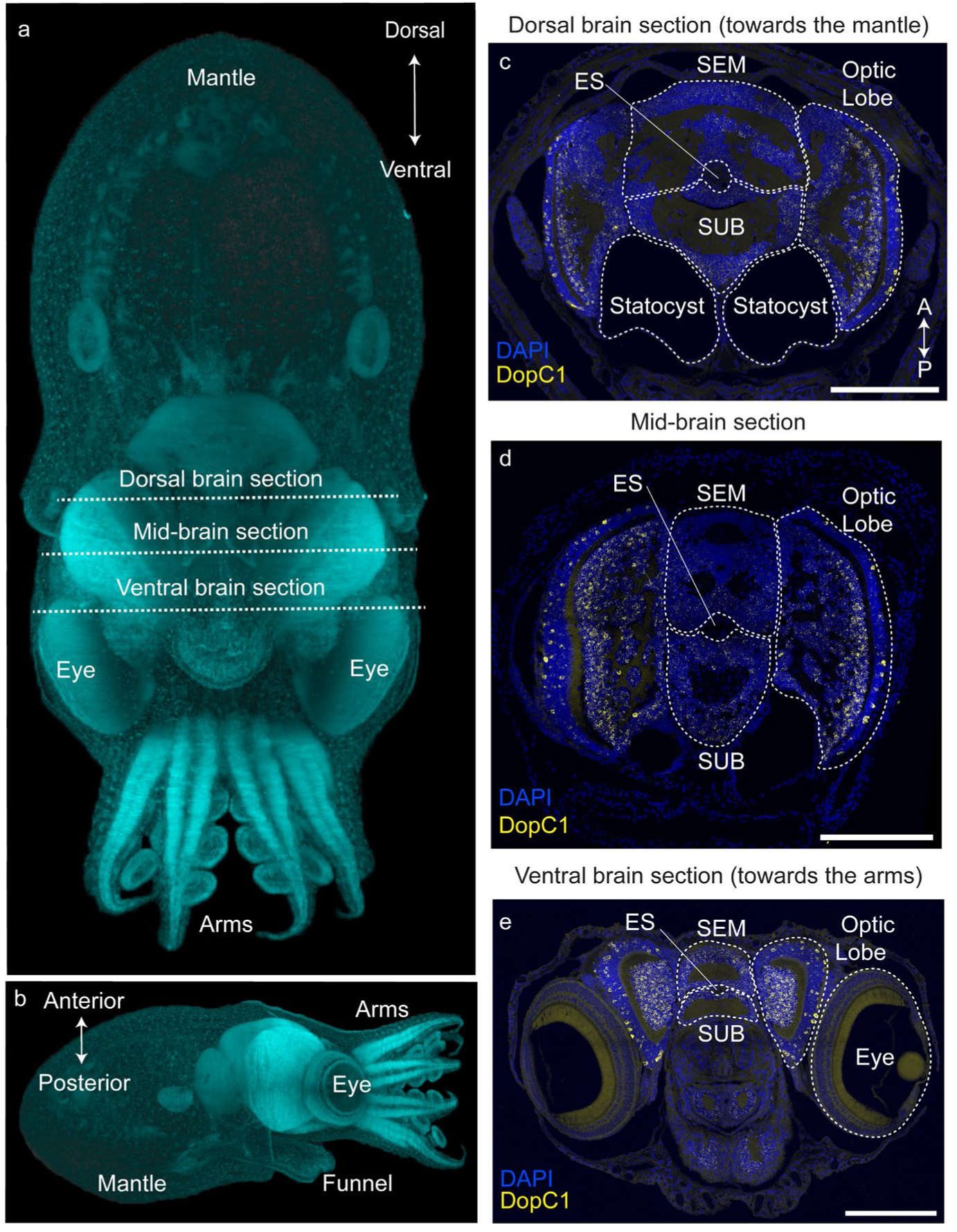
DopC1 expression in the central brain of *O. vulgaris* paralarvae. (**A** and **B**) Maximum intensity projection of DAPI labelled 1-dph *O. vulgaris* paralarva imaged with light sheet microscopy, reproduced with permission from Deryckere et al.^53^. (**C-E**) Representative HCR against DopC1 in yellow and nuclei (DAPI) in blue, at a dorsal position of the brain (towards the mantle) (**C**), at a mid position of the brain (**D**) and at a ventral position of the brain (towards the arms) (**E**). Scale bar: 200µm, SEM: supraesophageal mass of the central brain, SUB: subesophageal mass of the central brain, ES: oesophagus, A: anterior, P: posterior.

**Supp Fig. 8.**
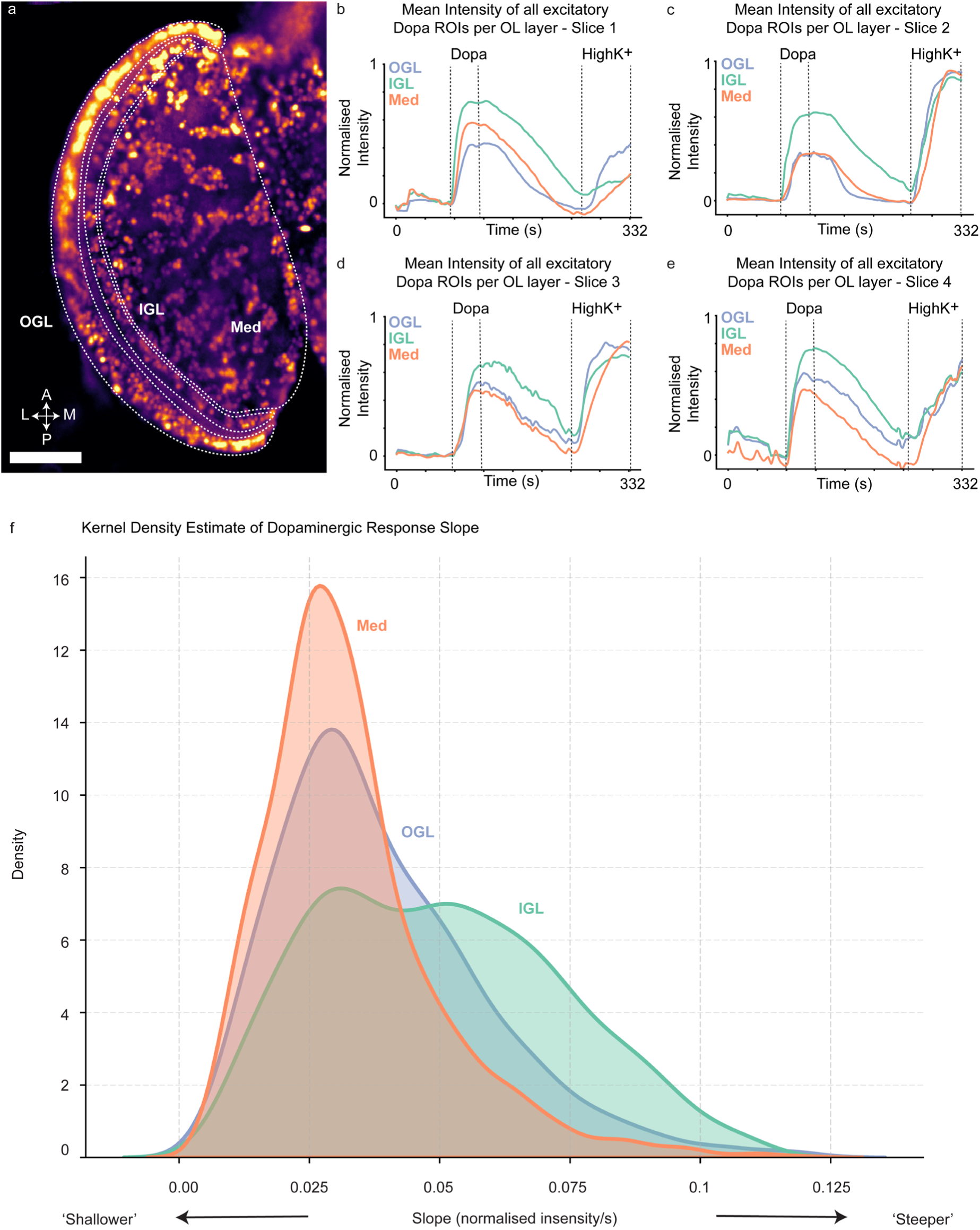
Additional analysis of calcium imaging on acute brain slices from *O. vulgaris* paralarvae during application of dopamine. Calcium imaging was performed on acute brain slices from 1-dph *O. vulgaris* paralarvae. (**A**) Maximum intensity projection of a representative brain slice used for calcium imaging. (**B-E)** The mean intensity over time for all ROIs in each optic lobe layer that were categorised as having an excitatory dopaminergic response are plotted as purple (OGL), green (IGL) and orange (Med) lines. Four different slices have been plotted. (**F**) Kernel density estimate of dopaminergic response slope per optic lobe layer. Scale bar: 100 µm, Dopa: dopamine, HighK^+^: high potassium solution, OGL: outer granular layer, IGL: inner granular layer, Med: medulla, A: anterior, P: posterior, M: medial, L: lateral.

**Supp Fig. 9.**
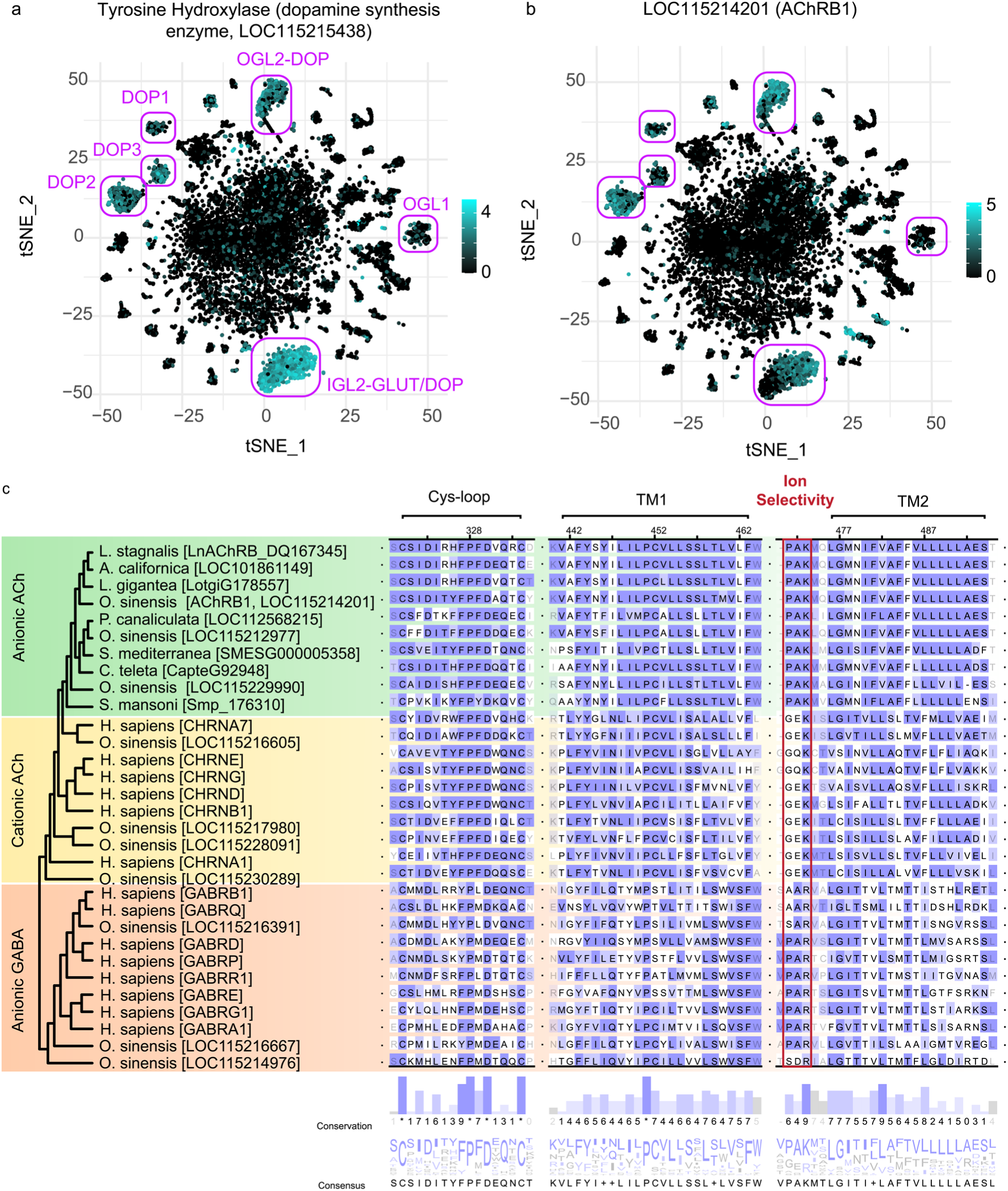
Additional expression, and sequence analysis of AChRB1 from *O. vulgaris*. tSNE of scRNA-seq of the *O. vulgaris* paralarval brain from^9^, showing the expression of dopaminergic marker th (tyrosine hydroxylase, LOC115215438) (**A**) and AChRB1 (LOC115214201) (**B**). Dopaminergic clusters are highlighted in magenta. Trimmed amino acid alignment showing a subset of anionic ACh, cationic ACh and anionic GABA LGCs highlighting multiple functional domains including the cys-loop region, transmembrane domain 1, the ion selectivity region and transmembrane domain 2 (**C**). Amino acid consensus and conservation values are shown below. Purple shading indicates BLOSUM62 scores indicating evolutionary divergence of individual sites. TM: transmembrane,

**Supp Fig. 10.**
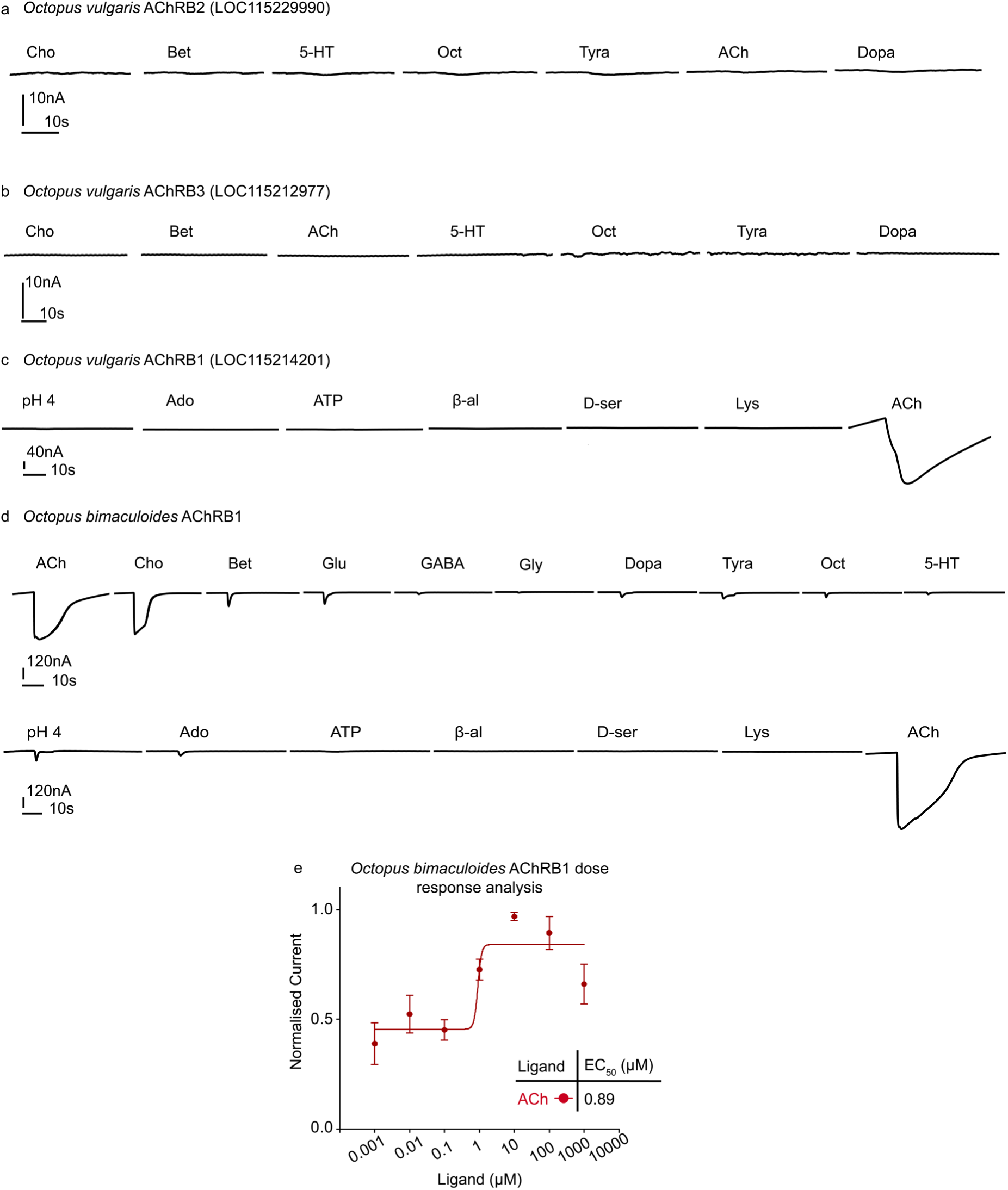
Functional characterisation of AChRB orthologs from *O. vulgaris* and *O. bimaculoides*. Representative TEVC traces from *Xenopus* oocytes expressing *O. vulgaris* AChRB2 (**A**), *O. vulgaris* AChRB3 (**B**), *O. vulgaris* AChRB1 (**C**), and *O. bimaculodes* AChRB1(**D**) during application of all candidate ligands. For D the top and bottom traces were from separate oocytes. (**E**) Dose response analysis curves for *O. bimaculoides* AChRB1 in response to ACh. This was acquired by performing TEVC recordings on *O. bimaculoides AChRB1* expressed in *Xenopus* oocytes from low to high concentrations. The adjacent table indicates the EC_50_ values for ACh in µM. Mean and standard error of the mean are plotted for 8 separate oocytes per ligand. The current was normalised to the max current in each oocyte. ACh: acetylcholine, Chol: choline, Bet: betaine, Glut: glutamate, Gly: glycine, Ado: adenosine, ATP: adenosine triphosphate, β-al: β-alanine, Lys: lysine, Dop: dopamine, Norepi: norepinephrie, Epi: epinephrine, Octo: octopamine, Tyra: tyramine, 5-HT: serotonin.

**Supp Fig. 11.**
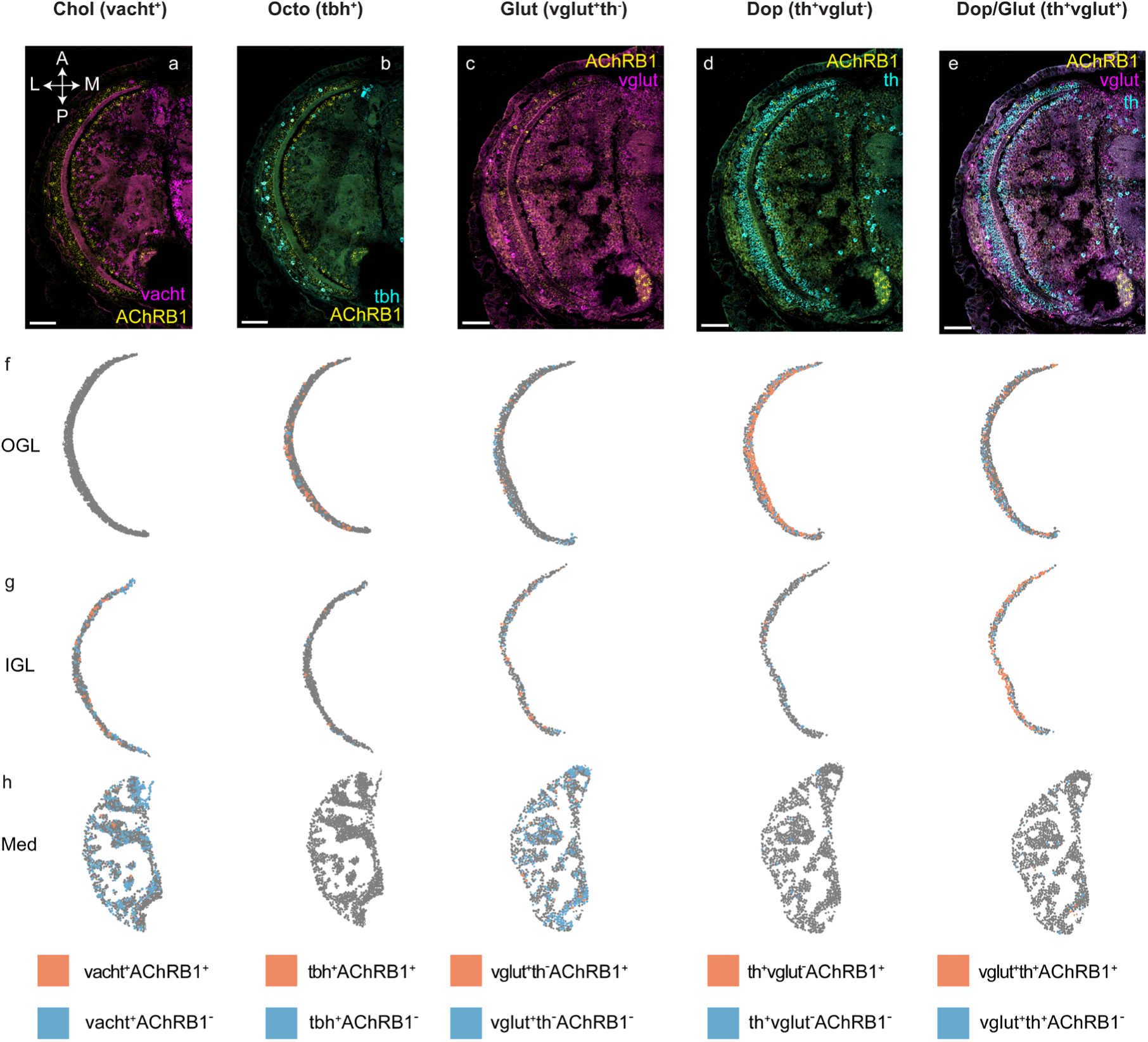
Quantitative and spatial analysis of AChRB1 expression in the brain of *O. vulgaris*. HCRs were performed on 1-dph *O. vulgaris* head transversal sections. Representative multiplex HCR against AChRB in yellow and vacht (cholinergic marker) in magenta (**A**), tbh (octopaminergic marker) in cyan (**B**), vglut (glutamatergic marker) in magenta (**C**) and th (dopaminergic marker) in cyan (**D**). (**C** and **E**) Representative multiplex HCR against AChRB in yellow, vglut (glutamatergic marker) in magenta (**C**) and th (dopaminergic marker) in cyan (**E**). (**F-H**) Representative HCR expression maps of Chol, Octo, Glut, Dop and Dop/Glut cell types in the OGL (**F**), IGL (**G**) and Med (**H**) of the OL, with cells positive for AChRB1 in orange) and those negative for AChRB1 in blue. The expression maps were quantified from the HCRs shown above and the results are summarised in Fig. 6 k. The orientation of all HCRs is the same. Cell type, layer, and gene names are as described in previous figures. Scale bar: 50 µm.

**Supp Fig. 12.**
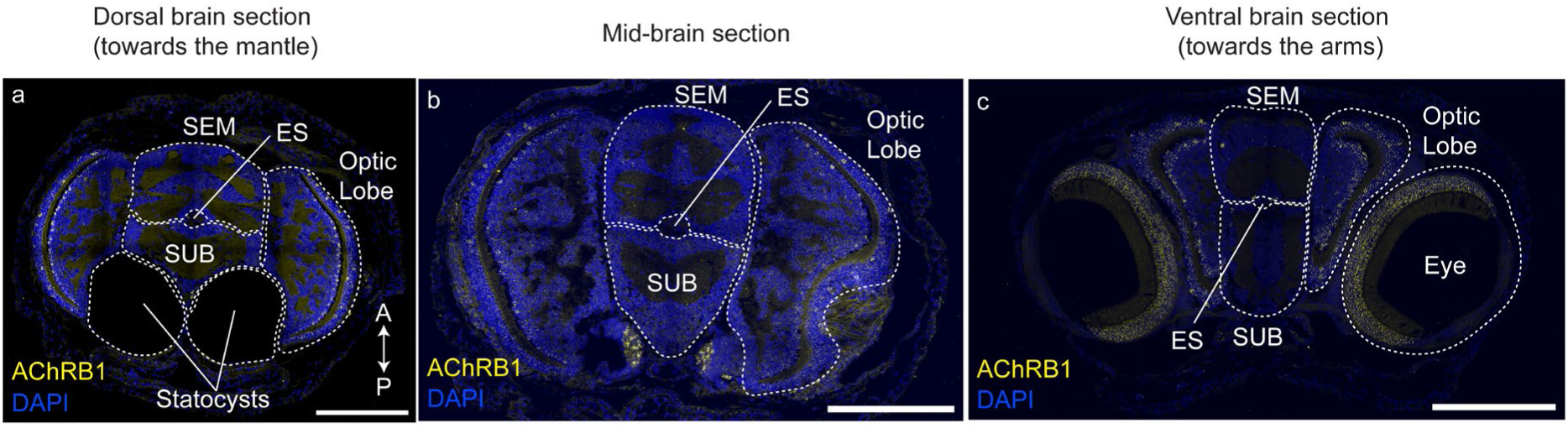
AChRB1 expression in the central brain of *O. vulgaris* paralarvae. Representative HCR against AChRB1 in yellow and nuclei (DAPI) in blue, at a dorsal position of the brain (towards the mantle) (**A**), at a mid-brain position (**B**) and at a ventral brain position (towards the arms) (**C**). For more details on ventral/dorsal orientations of the paralarvae see Supplemental Fig. 8. Scale bars: 200µm, SEM: supraesophageal mass of the central brain, SUB: subesophageal mass of the central brain, ES: oesophagus.

**Supp Fig. 13.**
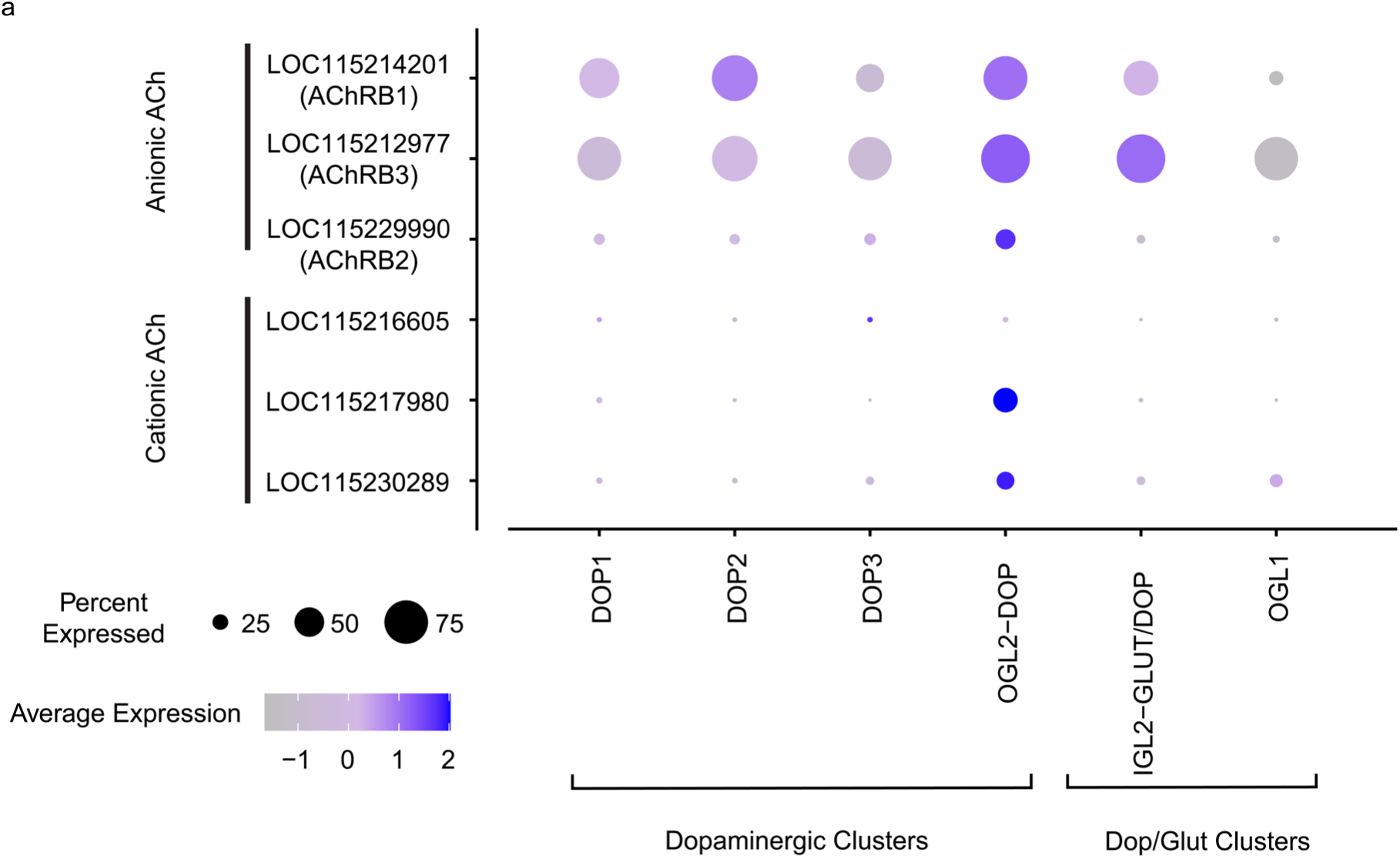
Expression analysis of anionic and cationic ACh-gated channels in dopaminergic and Dop/Glut cells. Dot plot of exclusively dopaminergic and Dop/Glut clusters from the scRNA-seq of the *O. vulgaris* paralarval brain^9^, showing the expression of anionic and cationic ACh genes. The size of the dots relates to the percent of cells in that cluster that express the gene, the heat map corresponds to the average expression of the gene in all cells in the cluster.

**Supp Fig. 14.**
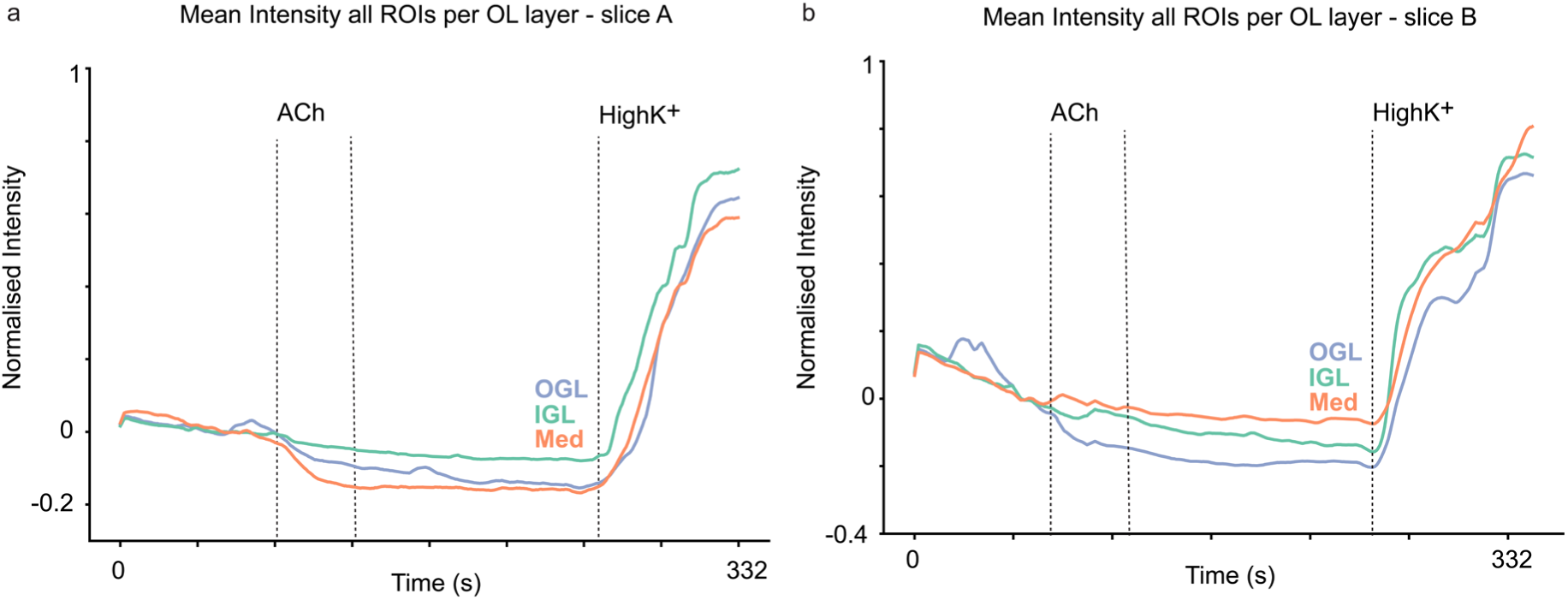
Additional analysis of calcium imaging on acute brain slices from O. vulgaris paralarvae during application of Ach. Calcium imaging was performed on acute brain slices from 1-dph *O. vulgaris* paralarvae during application of 100 µM ACh and HighK^+^. The mean intensity over time for all ROIs in each optic lobe layer are plotted as purple (OGL), green (IGL) and orange (Med) lines. Two different slices have been plotted (**A** and **B**). HighK^+^: high potassium solution, ACh: acetylcholine, OGL: outer granular layer, IGL: inner granular layer, Med: medulla.

**Supp Fig. 15.**
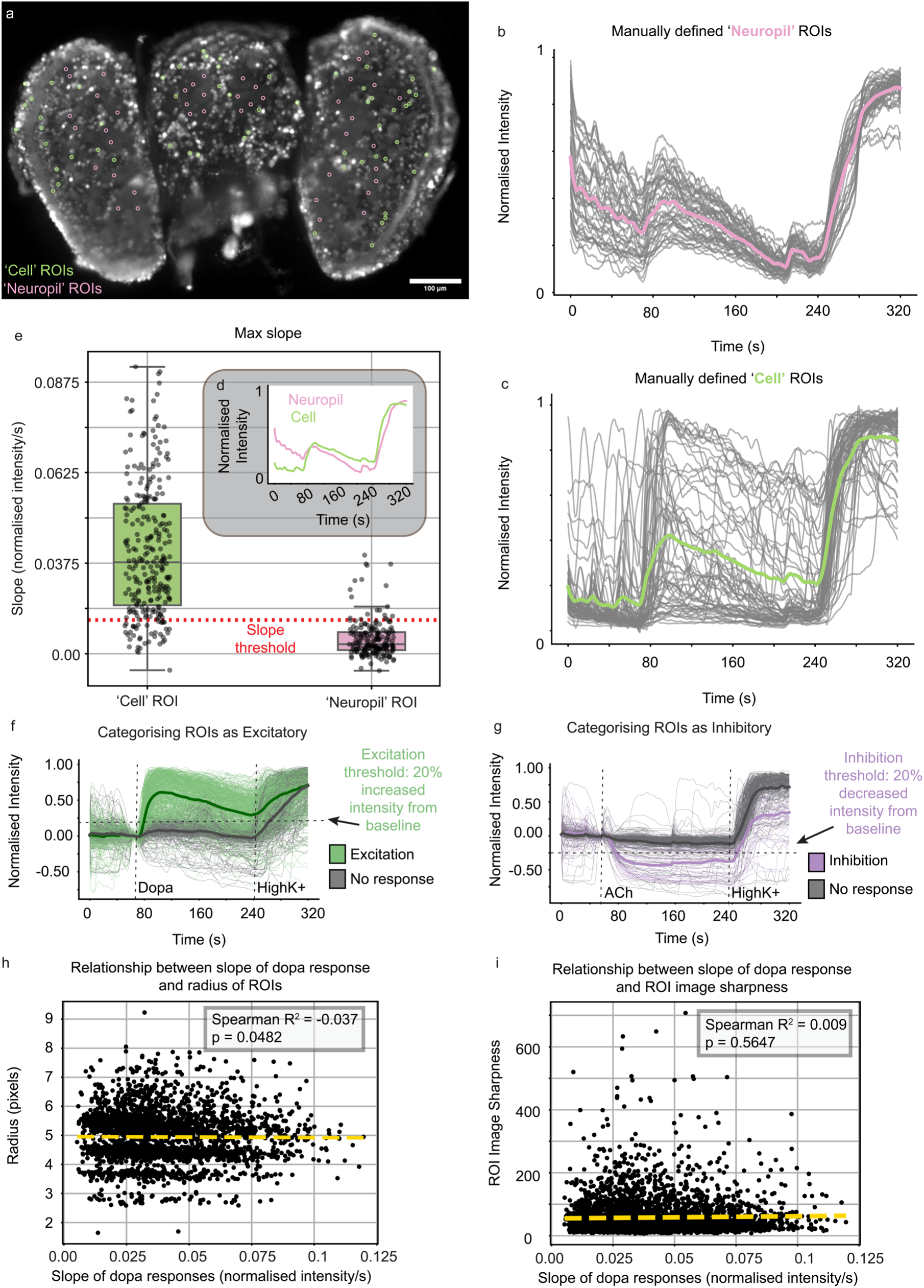
Calcium imaging methodology. Calcium imaging was performed on acute brain slices from 1-dph *O. vulgaris* paralarvae. Maximum intensity projection of a representative brain slice used for calcium imaging with manually defined ROIs for ‘Neuropil’ (pink) and ‘Cells’ (green) (**A**). Normalised mean intensity over time for all manually defined ‘Neuropil’ ROIs from A in grey. Mean trace is represented in pink (**B**). Normalised mean intensity over time for all manually defined ‘Cell’ ROIs from A in grey. Mean trace is represented in green (**C**). Mean ‘Neuropil’ and ‘Cell’ ROIs from B and C overlayed (**D**) The max slope was calculated for all manually defined ‘Cell’ and ‘Neuropil’ ROIs from five slices from five different paralarvae used for calcium imaging (**E**). The red dotted line is the mean + standard deviation of the ‘Neuropil’ ROI max slope. The normalised intensity of all traces from a representative slice that was exposed to 100 µM dopamine and HighK^+^ (**F**). The traces that were categorised as ‘excitation’ are in green and the traces that were categorised as ‘no response’ are in grey. The mean trace for each group is highlighted as a thicker grey or green trace. The normalised intensity of all traces from a representative slice that was exposed to 100 µM ACh and HighK^+^ (**G**). The traces that were categorised as ‘inhibition’ are in purple and the traces that were categorised as ‘no response’ are in grey. The mean trace for each group is highlighted as a thicker grey or green trace. A scatter plot comparing the relationship between the radius of a cell ROI (**H**) or the sharpness of the image surrounding a ROI (**I**) and the slope of excitatory dopaminergic responses. The results of a Spearman correlation test are seen as a dotted yellow line. Scale bar: 100 µm, Dopa: dopamine, ACh: acetylcholine, HighK^+^: high potassium solution, ROI: region of interest.

## Supplemental files

Supplemental file 1 – names and gene IDs for all receptors functionally characterised in this study.

Supplemental file 2 – citations for all receptors included in the phylogenetic tree which had a functional annotation.

Supplemental file 3 – protein sequence alignment for ‘cys-loop’ LGCs from major metazoan phyla.

Supplemental file 4 – full phylogenetic tree for ‘cys-loop’ LGCs from major metazoan phyla. All receptors that have been previously functionally characterised have their ligand highlighted in yellow.

Supplemental file 5 – list of HCR probes used in this study.

Supplemental file 6 – stl file for 3D printed cylinder used as a temporary well during calcium imaging experiments.

## Acknowledgements

The authors gratefully acknowledge Denise Walker and all other past and present members of the Schafer and Seuntjens labs for technical assistance and helpful discussions. *Dapnia magna* were kindly provided by the labs of Robby Stoks and Luc De Meester from KU Leuven, Belgium. The authors thank Joshua Rosenthal from the Marine Biological Laboratory for kindly providing the *Octopus bimaculoides* AChRB1 *Xenopus* plasmid. The authors also wish to acknowledge Gáspár Jékely and Milena Marinković from the Living Systems Institute, University of Exeter, for their advice on the phylogenetic analysis. The authors also wish to thank the Arckens lab for kindly allowing us to use their equipment for the calcium imaging experiments. Jon Howe and the LMB microscopy facilities team offered state of the art imaging facilities and expertise which the authors were very grateful for. The authors are also grateful to Benjamin Jenkins from the University of Cambridge for his expert advice on phylogenetic analysis. We would also like to thank the LMB Scientific Computing team and Rory Bedford for assistance with using the computer clusters. The authors are also grateful to all members of the VisLab team, particularly Elfy Chiang. Finally, the authors wish to thank all support and administrative staff at the LMB and KU Leuven.

## Author Contributions

A.C., E.S., and W.R.S conceptualised the project and designed the experiments. A.C. carried out the phylogenetic analysis. E.A. supplied the *O. vulgaris* paralarvae. A.C., I.H., and J.G.R. performed the *Xenopus* oocyte electrophysiology experiments. R.S. and M.Las. prepared *O. vulgaris* and *D. magna* cDNA. A.C., R.S., L.G., M.Lan., and A.M.J. performed the HCR experiments and acquired the images. J.B. developed the HCR image analysis pipeline and A.C. carried out the analysis and visualisation. A.C., R.S., and M.V.D. optimised and carried out the calcium imaging experiments. A.C. and H.A.O. developed the calcium image analysis pipeline and A.C. carried out the analysis and visualisation. A.C., E.S., and W.R.S wrote the original manuscript, and all authors read and critically revised the manuscript. E.S. and W.R.S supervised the project and provided funding/infrastructure.

## Materials & Correspondence

Requests for further information and resources should be directed to the corresponding author, William Schafer.

