## Supplementary figures and images for "Novel dopaminergic neurotransmission in the *Octopus* visual system"

### Supplemental file 4

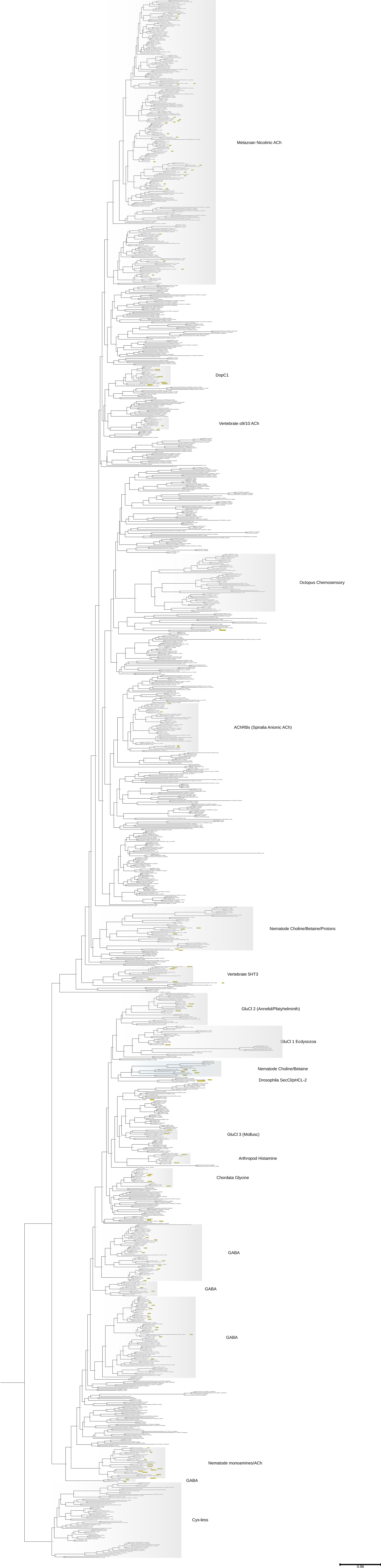
